# Targeted Genotyping of Variable Number Tandem Repeats with adVNTR

**DOI:** 10.1101/221754

**Authors:** Mehrdad Bakhtiari, Sharona Shleizer-Burko, Melissa Gymrek, Vikas Bansal, Vineet Bafna

## Abstract

Whole Genome Sequencing is increasingly used to identify Mendelian variants in clinical pipelines. These pipelines focus on single nucleotide variants (SNVs) and also structural variants, while ignoring more complex repeat sequence variants. We consider the problem of genotyping *Variable Number Tandem Repeats* (VNTRs), composed of inexact tandem duplications of short (6-100bp) repeating units. VNTRs span 3% of the human genome, are frequently present in coding regions, and have been implicated in multiple Mendelian disorders. While existing tools recognize VNTR carrying sequence, genotyping VNTRs (determining repeat unit count and sequence variation) from whole genome sequenced reads remains challenging. We describe a method, adVNTR, that uses Hidden Markov Models to model each VNTR, count repeat units, and detect sequence variation. adVNTR models can be developed for short-read (Illumina) and single molecule (PacBio) whole genome and exome sequencing, and show good results on multiple simulated and real data sets. adVNTR is available at https://github.com/mehrdadbakhtiari/adVNTR

## 1 Introduction

Next Generation Sequencing (NGS) is increasingly used to identify disease causing variants in clinical and diagnostic settings, but variant detection pipelines focus primarily on single nucleotide variants (SNVs) and small indels and to a lesser extent on structural variants. The human genome contains repeated sequences such as segmental duplications, short tandem repeats, and minisatellites which pose challenges for alignment and variant calling tools. Hence, these regions are typically ignored during analysis of NGS data. In particular, *tandem repeats* correspond to locations where a short DNA sequence or *Repeat Unit* (RU) is repeated in tandem multiple times. RUs of length less than 6bp are classified as Short Tandem Repeats (STRs), while longer RUs spanning potentially hundreds of nucleotides are denoted as *Variable Number Tandem Repeats* (VNTRs)(Shriver et al., 1993; Wright, 1994).

VNTRs span 3% of the human genome and are often found in coding regions where the repeat unit length is a multiple of 3 resulting in tandem repeats in the amino acid sequence. More than 1,200 VNTRs with a RU length of 10 or greater exist in the coding regions of the human genome(Tyner et al., 2016). Compared to STRs, which have been extensively studied (Gymrek et al., 2016; Ummat and Bashir, 2014; Liu et al., 2017; Willems et al., 2017; Dolzhenko et al., 2017), VNTRs have not received as much attention. Nevertheless, multiple studies have linked variation in VNTRs with Mendelian diseases (*e.g*., Medullary cystic kidney disease(Kirby et al., 2013), Myoclonus epilepsy(Lalioti et al., 1997), and FSHD(Lemmers et al., 2002)) and complex disorders such as bipolar disorder (Table 1). In some cases, the disease associated variants correspond to point mutations in the VNTR sequence (Kirby et al., 2013; Ræder et al., 2006) while in other cases, changes in the number of tandem repeats (RU count) show a statistical association (or causal relationship) with disease risk. For example, the insulin gene (INS) VNTR has an RU length of 14 bp with RU count varying from 26 to 200(Pugliese et al., 1997). Variation in this VNTR has been associated with expression of the INS gene and risk for type 1 diabetes (OR = 2.2) (Durinovic-Belló et al., 2010). Notwithstanding these examples, the advent of genome-wide SNP genotyping arrays led to VNTRs being largely ignored. They have been called ‘the forgotten polymorphisms’(Brookes, 2013).

**Table 1:**
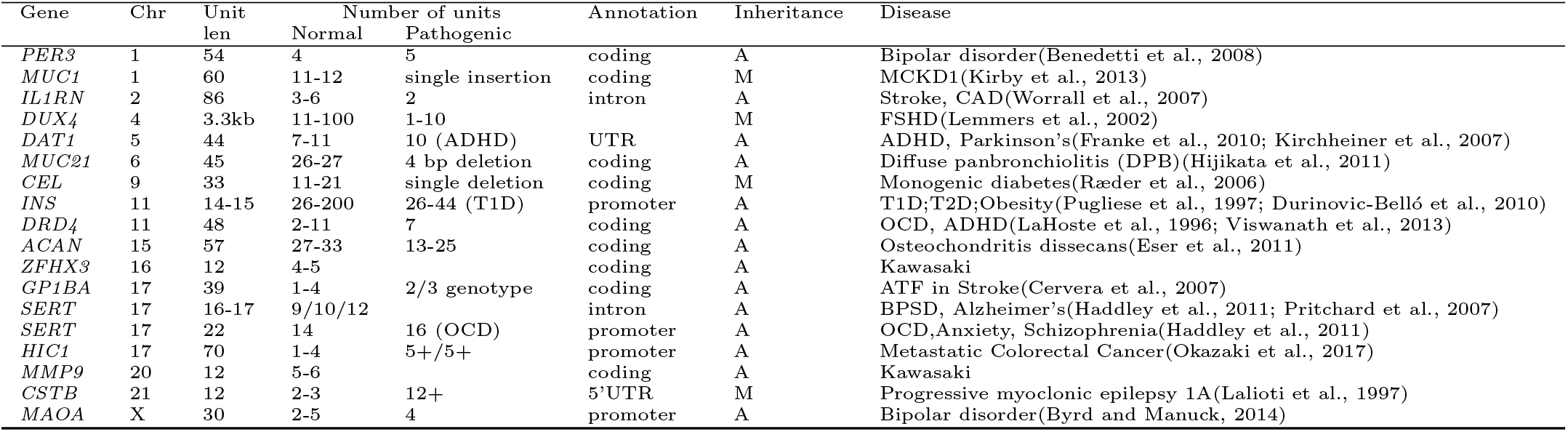
**Disease-linked VNTRs** are generally distinguished from STRs by a longer length (≥ 6) of the repeating unit. ‘M’ denotes Mendelian inheritance, while ‘A’ represents possibly complex inheritance captured via Association. As it is difficult to genotype VNTRs, most cases have been determined via association, but the inheritance mode could be high penetrance.

VNTRs were originally used as markers for linkage mapping since they are highly polymor phic with respect to the number of tandem repeats at a given VNTR locus(Gelfand et al., 2014). Traditionally, VNTR genotyping required labor intensive gel-based screens which limited the size of large population based studies of VNTRs (Orita et al., 1989). Whole genome sequencing has the potential to detect and genotype all types of genetic variation, including VNTRs. However, computational identification of variation in VNTRs from sequence remains challenging. Existing variant calling methods have been developed primarily to identify short sequence variants in unique DNA sequences that fall into a reference versus alternate allele framework, which is not well suited for detecting variation in VNTR sequences.

Genotyping VNTRs in a donor genome sequenced using short (Illumina) or longer single molecule reads, requires the following: (a) *recruitment of reads* containing the VNTR sequence; (b) *counting RUs* for each of the two haplotypes; (c) *identification of indels within VNTRs*; and (d) identification of mutations within the VNTR. Mapping tools such as BWA(Li and Durbin, 2009) and Bowtie2(Langmead and Salzberg, 2012) can work for read recruitment for STRs, but are challenged by insertion/deletion of larger repeat units. Mapping issues also confound existing variant callers, including realignment tools such as GATK IndelRealigner(DePristo et al., 2011) if the total VNTR length is larger than the read length. This is because reads contained within the VNTR sequence have multiple equally likely mappings and therefore will be mapped randomly to different locations with low mapping quality(Kirby et al., 2013). Detection of point mutations in long VNTRs requires integrating information across the entire VNTR sequence. For VNTRs whose total sequence length (RU count times the RU length) is much longer than the read length, detection of SNVs and indels is not feasible using existing variant callers. We focus mainly on problems (a,b) relating to recruitment and RU counting. For problem (c), we focus on difficult case of large (≥ 250bp) VNTRs within coding regions where the indel shifts the translation frame. We do not tackle problem (d) in this manuscript.

Other tools have addressed the problem of RU count estimation, focusing on the related problem of STR genotyping. Some of these tools do not accept large repeating patterns as input (Willems et al., 2017; Liu et al., 2017). Others require all repeat units to be near-identical(Dolzhenko et al., 2017; Ummat and Bashir, 2014). In particular, ExpansionHunter(Dolzhenko et al., 2017) looks for exact matches of short repeating sequence within flanking unique sequences, and works for STRs, but not as well with the larger VNTRs with variations in RUs (Results). VNTRseek(Gelfand et al., 2014) detects a VNTR-like pattern in reads and aligns it to tandem repeats, but uses a complex alignment process making it difficult to run the tool. Alignment based tools need to align reads at both unique ends, which may not be possible for short (Illumina) reads. Single molecule reads (*e.g*., PacBio(Eid et al., 2009), Nanopore(Clarke et al., 2009)) can span entire VNTR regions, but it is difficult to estimate the RU count directly since the distance between the flanking regions varies dramatically from read to read due to an excess of indel errors. For example, 14 reads spanning the SERT VNTR in the in the PacBio sequencing data of NA12878 individual from Genome in a Bottle(Zook et al., 2016) included fifteen distinct lengths between 292bp and 385bp, leading to length-based RU count estimates 13, 14, 15, 16, and 18 for the diploid genome.

In contrast to methods like VNTRseek which seek to *discover/identify* VNTRs, we describe a method, adVNTR, for *genotyping VNTRs* at targeted loci in a donor genome. For any target VNTR in a donor, adVNTR reports an estimate of RU counts and point mutations within the RUs. It trains Hidden Markov Models (HMMs) for each target VNTR locus, which provide the following advantages: (i) it is sufficient to match any portions of the unique flanking regions for read alignment; (ii) it is easier to separate homopolymer runs from other indels helping with frameshift detection, and to estimate RU counts even in the presence of indels; (iii) each VNTR can be modeled individually, and complex models can be constructed for VNTRs with complex structure, along with VNTR specific confidence scores. For longer VNTRs not spanned by short reads, adVNTR can still be used to detect indels, while providing lower bounds on RU counts. Also, exact estimates for RU counts could be made for shorter VNTRs. Using simulated data as well as whole-genome sequence data for a number of human individuals, we demonstrate the power of adVNTR to genotype VNTR loci in the human genome.

## 2 Results

Our method, adVNTR, requires training of separate HMM models for each combination of target VNTR and sequencing technologies. The detailed training procedure is described in Methods. Given trained models, adVNTR genotypes the VNTRs in three stages: (i) Selection of reads that contain VNTR locus (read recruitment); (ii) RU count estimation; and, (iii) variant detection. We report results on performance of adVNTR in each of these stages using simulated and read datasets based on short-read (Illumina) and single molecule (PacBio) technologies.

### HMM training

Initial HMMs were trained using multiple alignments of RU sequences from the reference assembly hg19(Lander et al., 2001), as described in methods. Similarly, HMMs were trained for the left flanking and right flanking regions for each VNTR. The HMM models were augmented using data from Genome in a Bottle (GIAB) project (NA12878 WGS). VNTR models were trained for VNTRs in coding and promoter regions of the genome, for both Illumina (1755 models) and PacBio (2944 models; Supplementary Material “Selecting Target VNTRs”). Subsequently, we tested performance for (a) read-recruitment, (b) counting of Repeat Units, and (c) detection of indels.

### Test Data

To evaluate performance for *PacBio*, we simulated haplotypes for each of the 2944 VNTRs, revising the RU count to be ±3 of the RU count in hg19, and setting 1 as the minimum RU count. We simulated haplotype reads (15X coverage) using SimLoRD(Stöcker et al., 2016) and aligned those reads to hg19 using Blasr(Chaisson and Tesler, 2012). For Illumina sequencing, we used ART(Huang et al., 2011) to simulate haplotype WGS (shotgun 150bp) reads at 15X coverage for each VNTR and simulated VNTR haplotype with changes in RU counts similar to PacBio. Pairs of haplotypes were merged to get (30X coverage) diploid samples. The resulting data-sets were called *PacBioSim* and *IlluminaSim*, respectively (Supplementary Material “Test Datasets”). To evaluate performance of frameshift identification, we collected a set of 115 VNTRs (Supplementary Material “Selecting Target VNTRs”). For each VNTR, we simulated haplotypes that contain a deletion or an insertion in the VNTR (Supplementary Material “Test Datasets”). We simulated reads from each of these haplotypes and merged pairs of halpotypes to obtain diploid samples. We denote this data-set as *IlluminaFrameshift*.

### Read recruitment

adVNTR takes a collection of VNTR models as input, and as a first step, recruits reads that map to any of the VNTRs in the list. In testing recruitment for PacBio, we found that alignment tools such as Blasr perform well in recruiting VNTR reads even in the presence of deletions and insertions (data not shown) and used Blasr for all read recruitment. For *Illumina* reads, we tested adVNTR read-recruitment for all 1775 VNTRs using IlluminaSim, and compared against mapping tools BWA-MEM, Bowtie2, and BLAST. adVNTR achieves much greater recall while maintaining or exceeding the precision of other tools (Fig. 1 and Fig. S3). Specifically, adVNTR recall was 100% for 99.9% of the VNTRs, whereas the next best tool (BWA-MEM) achieved this only for 68.2% of the VNTRs. The other mapping tools lose mapping sensitivity when RU counts are increased or decreased (large indels), and perform best when the RU counts are the same as reference (Fig. S2A-C), partially explaining their lower recall.

**Figure 1:**
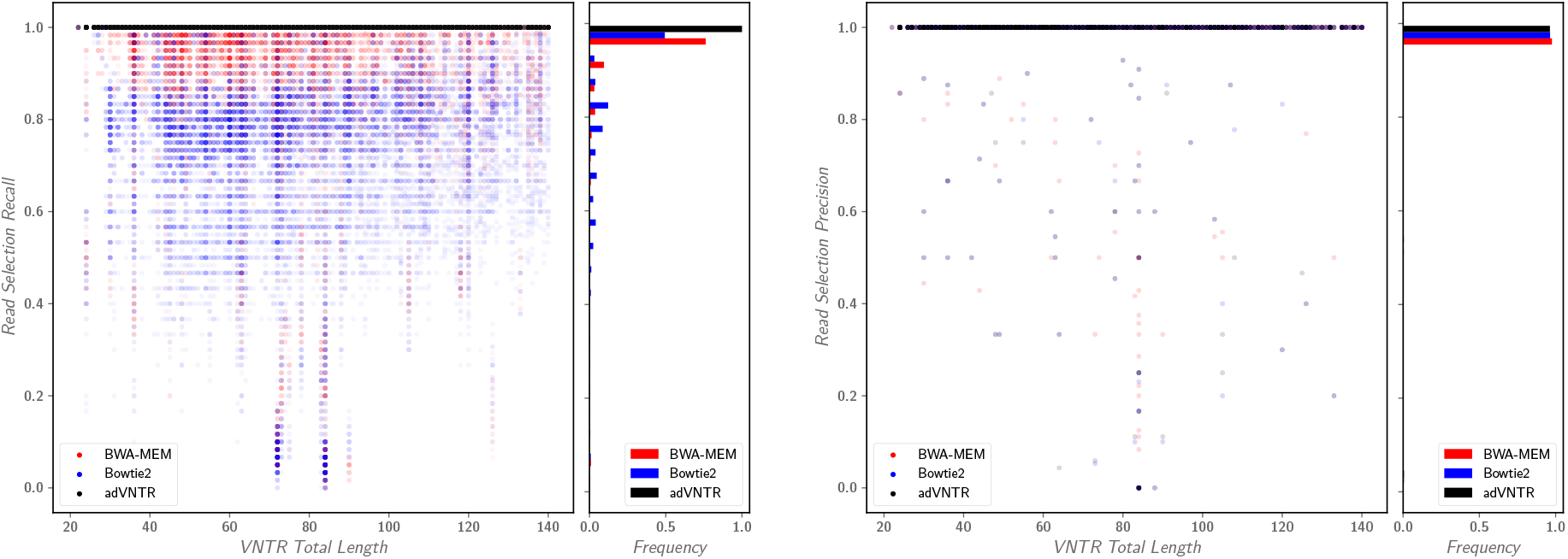
Read recruitment quality on Illumina reads. (A) Comparison of the recall (# true recruited reads/# true reads) of adVNTR read recruitment against BWA-MEM and Bowtie2, as a function of VNTR length for 1775 VNTRs with different counts (31, 788 tests). Each dot corresponds to a separate test. (B) Precision (# true recruited reads/# recruited reads) of read recruitment.

### VNTR genotyping (RU count estimation) with PacBio reads

Recall that sequencing (particularly homopolymer) errors can cause lengths to change, particularly for short RU lengths and larger RU counts. To test adVNTR performance on PacBioSim, we compared against a naïve method that estimates RU counts based on read length between the flanking regions from the consensus of reads that cover VNTR. Detailed performance on three exemplars (INS, CSTB, and HIC1) gene showed high genotype accuracy for adVNTR over a wide range of RU counts, and coverage (Fig. 2A. Similar results were obtained for all 2944 VNTRs (Fig. 2B). Overall, 98.45% of adVNTR estimates were correct while 26.45% of estimates made by naïve method were correct. As it is difficult for the naïve mthod to call heterozygotes, we also compared on the subset of test data with homozygous RU counts. 97.95% of adVNTR estimates were correct, while the consensus method was correct in 66.16% of samples (Fig. S4). adVNTR estimates were uniformly good except at low sequence coverage. To test for accuracy with changing RU counts, we simulated different RU counts for individuals at 3 VNTRs (Table S4). adVNTR RU counts showed 100% accuracy in each of the 52 different samples tested.

**Figure 2:**
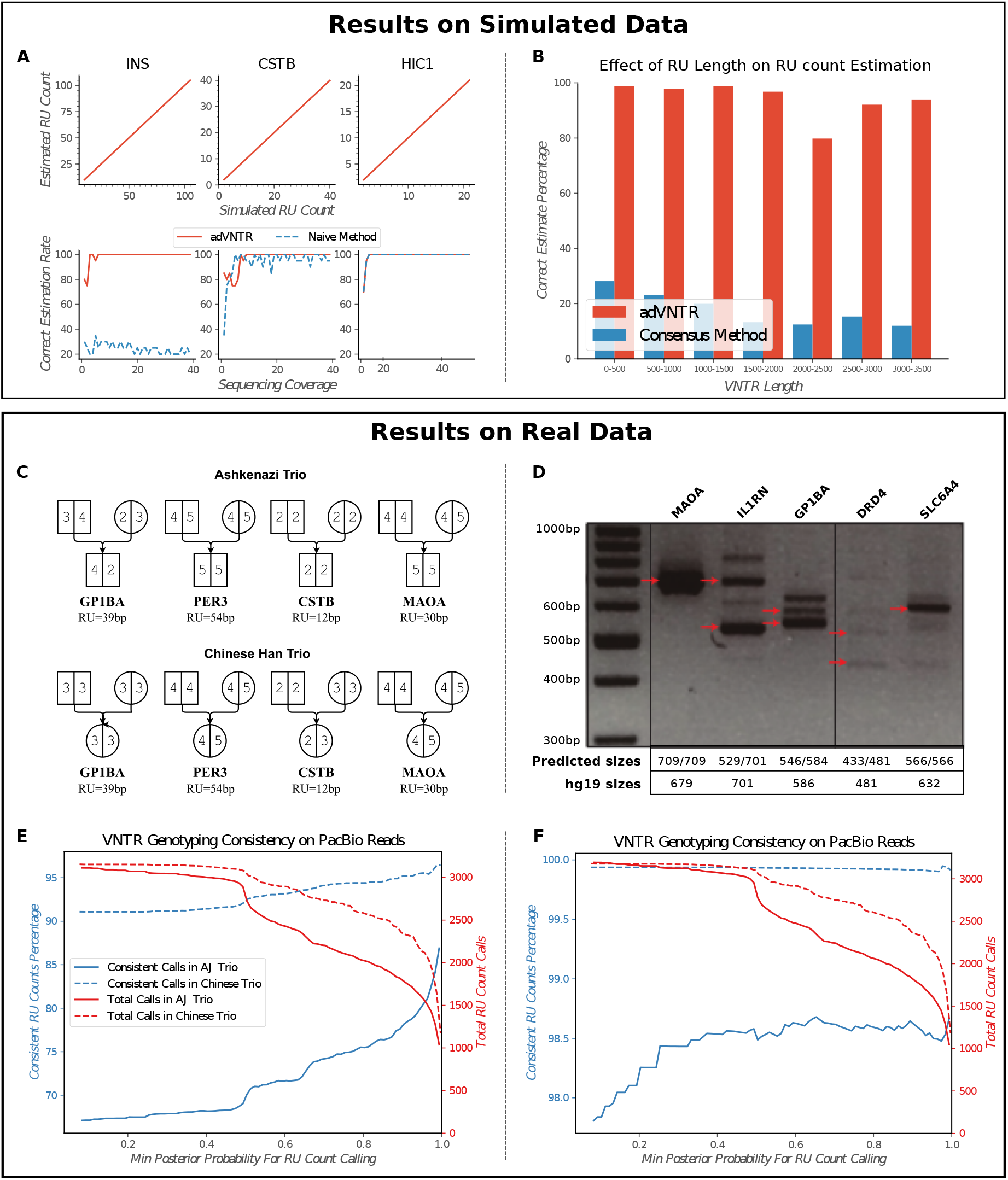
VNTR genotyping using PacBio data. (A) RU count estimation on simulated PacBio reads as a function of RU count and coverage for 3 medically relevant VNTRs: INS (RU length 14bp), CSTB (12 bp), and HIC1 (70bp). adVNTR performance is compared to a naïve method. (B) The effect of RU length on count accuracy over 2944 VNTRs (30418 tests). (C) Mendelian consistency of genotypes at 4 VNTR loci in the Chinese Han and Ashkenazi trios. Note that MAOA results are consistent with its location on Chr X. (D) LR-PCR based validation of genotypes at 5 disease-linked VNTRs in NA12878. Red arrow correspond to VNTR lengths estimated by multiplying predicted RU counts with RU lengths. (E) Fraction of consistent calls and number of calls across 2944 VNTRs in AJ and Chinese trios from GIAB and NCBI-SRA. (F) Fraction of consistent calls allowing for off-by-one errors.

To test performance on real data where the true VNTR genotype was not known, we checked for Mendelian inheritance consistency in the AJ trio from Genome in a Bottle (GIAB)(Zook et al., 2016) and a Chinese Han trio from NCBI SRA (accession PRJEB12236). On four disease related VNTRs, adVNTR predictions were consistent in each case (Fig. 2C). On the 2944 genic VNTRs, the trio consistency of adVNTR calls was correlated with coverage. At a posterior probability threshold of 0.99, 86.98% of the calls in the AJ trio, and 97.08% of the calls in the Chinese trio, were consistent with Mendelian inheritance (Fig.2E). Many of the discrepancies could be attributed to low coverage and missing data. Increasing sequence coverage threshold from 5× to 10 × increased the average posterior probability from 0.91 to 0.98 and resulted in improved RU count accuracy (Fig. S5). Also, many of these discrepancies in RU counts were off-by-one errors (Fig. S6). These off-by-one discrepancies could be acceptable for Mendelian disease testing as the pathogenic cases often have large changes in RU counts. Treating the off by one counts as correct, we found that 98.66% and 99.91% of the high confidence calls in AJ and Chinese trios, respectively, were consistent (Fig.2F). Finally, some of the off-by-one counts could be natural genetic variation.

We also performed a long range (LR)PCR experiment on the individual NA12878 to assess the accuracy of the adVNTR genotypes using PacBio data (Table S2 and Table S3). The observed PCR product lengths (black bands in Fig. 2D) were consistent with the adVNTR predictions (red arrows), while being different from the hg19 reference RU count. adVNTR correctly predicted all VNTRs to be heterozygous with the exception of SLC6A4, that was predicted to be homozygous.

While we could not get the VNTR discovery tool VNTRseek(Gelfand et al., 2014) to run on our machine (personal communication), we observed that the authors had predicted 125 VNTRs in the Watson sequenced genome(Wheeler et al., 2008), and 75 VNTRs in two trios as being polymorphic. In contrast, analysis of the PacBio sequencing data identified >500 examples of polymorphic VNTRs that overlap with coding regions. The results suggest that variation in RU counts of VNTRs and their role in influencing phenotypes might be greater than previously estimated.

### RU counting with Illumina

The adVNTR estimate correctly matched both RU counts in 91.6% of the cases in the IlluminaSim dataset (1775 VNTRs with up to 21 diploid RU counts each) and matched at least one RU count in 97% of the cases (Fig. 3A,B). Most of the discrepancies occurred in VNTRs with longer lengths not covered by Illumina reads (Fig. 3C,D). While there was a drop in accuracy for increasing lengths, 84% of the genic VNTRs are shorter than 150bp, and could be genotyped with 94.6% accuracy. Tools such as VNTRseek require at least 20bp flanking each side of the VNTR and do not return a result for VNTRs with total length greater than 110bp, while adVNTR could predict the genotype correctly in a majority of those cases (Supplementary Material “VNTRseek”). ExpansionHunter, a tool designed primarily for STR genotyping (Dolzhenko et al., 2017) provided incorrect estimates in over 90% cases from this data-set (Fig. S7). ExpansionHunter makes the assumption that the different RUs are mostly identical in sequence which is valid for STRs but not for most VNTRs, and we tested this through 52 samples on three VNTRs. adVNTR predicted the correct genotype in all but 6 cases, with erroneous calls only in the case of high RU counts where the read length did not span the VNTR perfectly, while ExpansionHunter did not return the correct estimate in most cases (Table S4).

**Figure 3:**
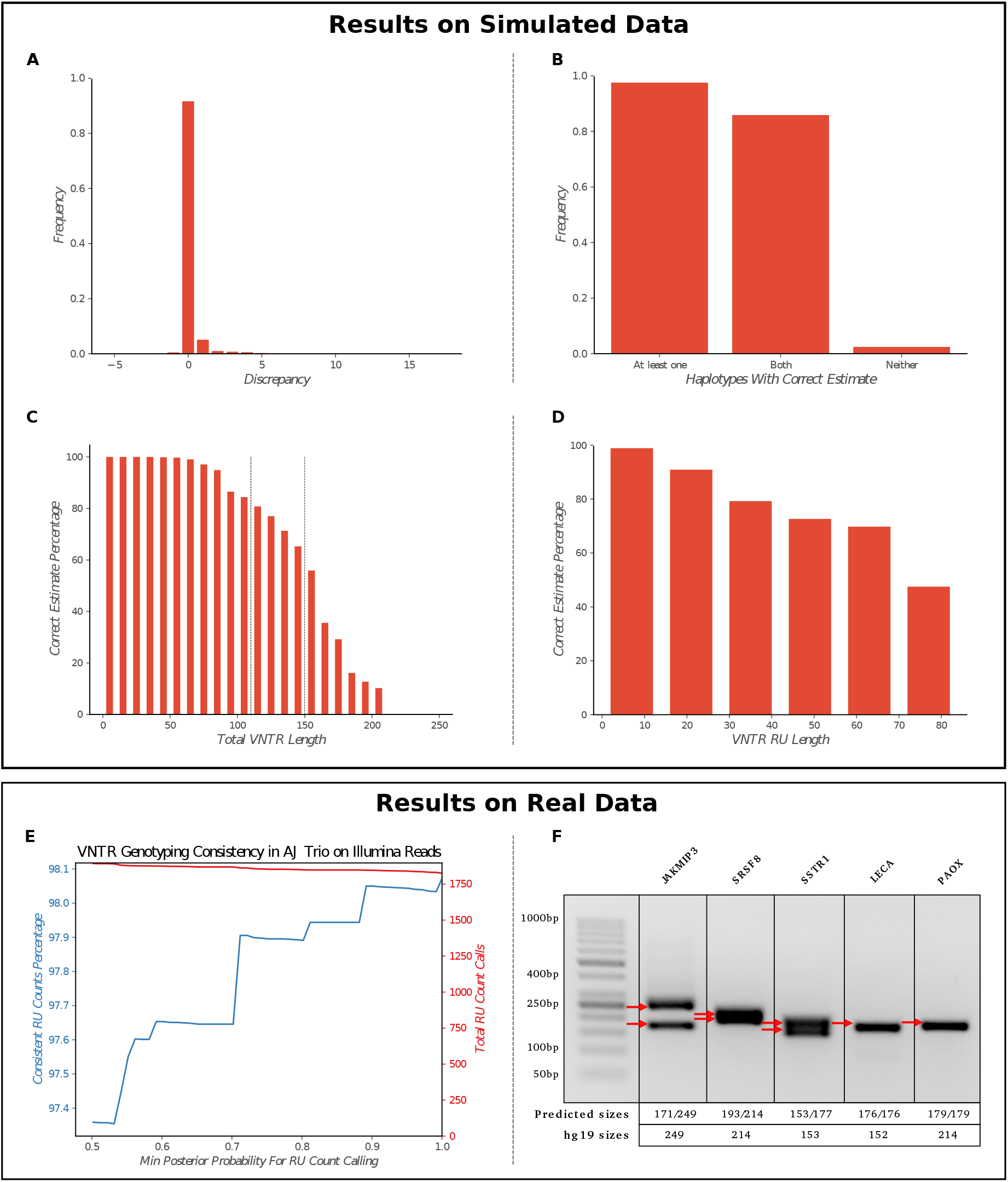
VNTR genotyping using Illumina data. (A-D) Correctness of RU count prediction for 1775 coding VNTRs in the IlluminaSim dataset, described by (A) RU count dscrepancy, (B) haplotypes with correct estimates, (C) correctness as a function of VNTR length, and (D) RU length. (E) Consistency of adVNTR calls on the AJ trio WGS data from GIAB. Red line describes the cumulative number of calls made at specific posterior probability cut-offs. (F) Gel electrophoresis based validation of adVNTR calls on 5 short VNTRs using WGS of individual NA12878 from GIAB. Red arrows correspond to VNTR lengths estimated by multiplying the RU lengths with the estimated RU counts.

On the AJ trio from GIAB, 98.08% of the high confidence adVNTR calls were consistent with Mendelian inheritance (Fig. 3E). Note that 95.93% of all calls were high confidence (posterior probability ≥ 0.99). We validated adVNTR calls on 12 VNTRs using Gel electrophoresis (Table S3). adVNTR predicted the correct RU counts in all cases, except in two cases where the PCR primers failed to produce a band (Fig. 3F, S8). We also compared adVNTR against ExpansionHunter on 7 disease related short VNTRs in the AJ trio and obtained similar results (Table S5).

To test adVNTR for population-scale studies of VNTR genotypes using WGS data replacing labor intensive gel electrophoresis(Byrd and Manuck, 2014; Cervera et al., 2007), we scanned the PCR-free WGS data for 150 individuals (50 in each population) obtained from 1000 genomes project(Consortium, 2015). We observed population specific RU counts (frequency difference > 10%) in 97 of 202 VNTRs tested (Table S7). Fig. 4 shows the RU count frequencies for a disease-linked VNTR in the coding region of CSTB and a coding VNTR in CCDC66. The results suggest an increase in VNTRs with higher RU counts with an increase in divergence time from Africa. Thus RU3 is more prevalent in both VNTRs. We also observed RU4 in CSTB6 VNTR in the Asian and European populations, where RU counts 4 and above have been associated with progressive myocolonal epilepsy (Lalioti et al., 1997).

**Figure 4:**
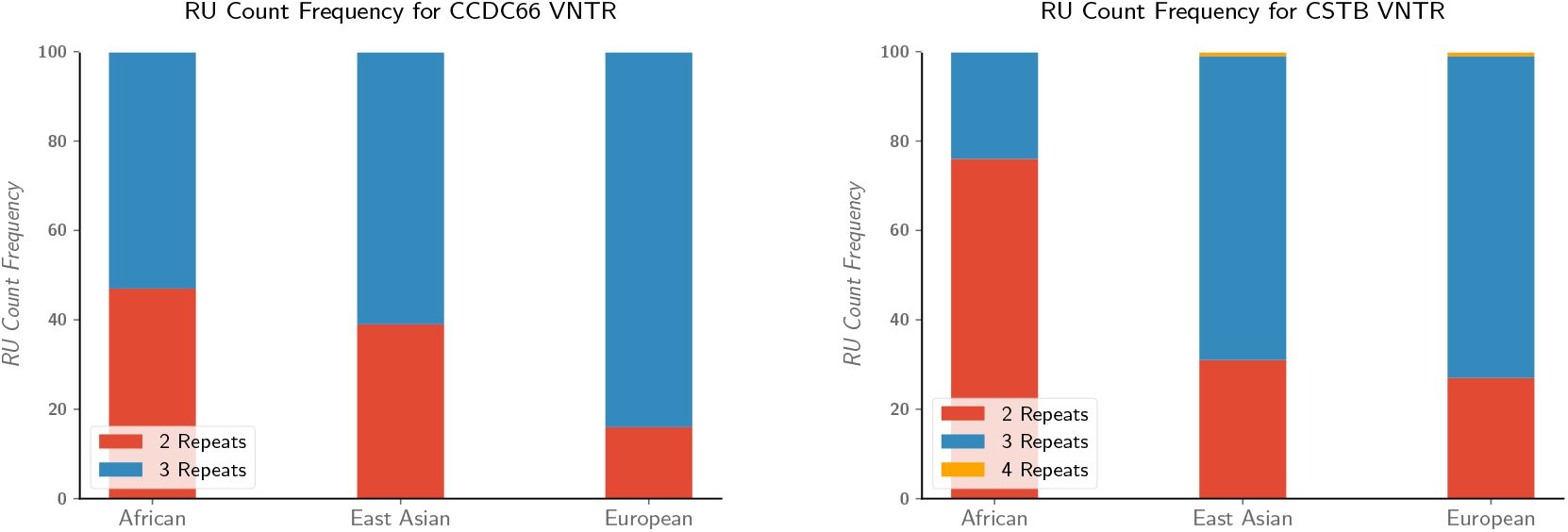
Population-scale genotyping of VNTRs. (A) RU count frequencies for the VNTR in CCDC66 gene, and (B) CSTB in African, Asian, and European population samples from 1000 genomes project. RU counts of 4 and higher in CSTB are associated with myoclonal epilepsy.

### VNTR mutation/indel detection

As a proof of concept for other applications, we tested indel detection, focusing in particular on frameshifts in coding VNTRs. The CEL gene is known to contain a VNTR where a deletion changes the coding frame. We simulated Illumina reads from 20 whole genomes after introducing a single insertion or deletion in the middle of the VNTR region in the CEL gene. As a negative control, we simulated 10 WGS experiments with a range of sequence coverage values. We ran adVNTR, Samtools mpileup(Li, 2011), and GATK HaplotypeCaller(DePristo et al., 2011) which uses GATK IndelRealigner, to identify frameshifts in each of the simulated datasets, and the 10 control datasets. On the control data, none of the tools found any variant. On the simulated indels, adVNTR made the correct prediction in each case (Suppl. Table S6), while Samtools and GATK were unable to predict a single insertion or deletion. This result is not surprising as the reads have poor alignment scores, and the indel can be mapped to multiple locations (Suppl. Fig. S9)(Robinson et al., 2011). We note that mapping ambiguity in aligning each read made it difficult to pinpoint the location of single indel. However, by integrating the information across all reads, we could predict the occrrence of a frameshift in the VNTR. We next tested adVNTR frameshift prediction on the 115 VNTRs in the IlluminaFrameshift dataset, simulating 4090 total cases. Overall, the frameshifts in the VNTR regions were predicted with 51.7% sensitivity and 86.8% specificity, in contrast with the 49.7, 43.5% sensitivity, specificity achieved by GATK. Detailed performance of methods for each VNTR is available in Table S7. Note that the performance is model specific and depends upon the similarity of different Repeat Units in a VNTR. For 29 of the 115 VNTRs, adVNTR showed high sensitivity (≥90%) and specificity (100%).

As frameshifts in the VNTR region of the CEL gene have been linked to a monogenic form of diabetes(Ræder et al., 2006), we tested for frameshifts in CEL using whole Exome sequencing (WES) data from 2,081 cases with Type 2 Diabetes (Fuchsberger et al., 2016) and compared the numbers to 2,090 control individuals. WES data analysis is challenging as high GC-content makes it difficult to PCR-amplify this VNTR. adVNTR found that while none of the controls had any evidence of a frameshift, 8 of the 2,081 diabetes cases showed a frameshift in this VNTR region (Suppl. Fig. S10).

### Compute requirements for genotyping

adVNTR is multi-threaded. In genotyping mapped PacBio reads at 30X coverage, adVNTR took 6 hours using Intel Xeon(R) 4-core CPUs (≤ 24 CPU-hours) to genotype all 2944 VNTRs, and 14:15 hours (≤ 57 CPU-hours) for 70X coverage. For Illumina reads at 40X coverage, adVNTR took 87:30 cpu-hours on a single core to complete read recruitment as well as genotyping of 1775 VNTRs.

## 3 Discussion

The problem of genotyping VNTRs (determining diploid RU counts and mutations) is increasingly important for clinical pipelines seeking to find the genetic mechanisms of Mendelian disorders. As VNTRs have not been extensively studied, existing research is often focused on their discovery. One of the contributions of this paper is the separation of initial VNTR discovery from VNTR genotyping, and a focus on the genotyping problem. adVNTR genotypes VNTRs using a hidden markov model for each target VNTR, providing a uniform training framework, but still allowing us to tailor the models for complex VNTRs on a case by case basis. The problem of mismapping due to indels introduced by changing RU counts confounds most mapping based tools, but is solved here by collapsing all RU copies and building HMMs that allow for variation in the RUs. adVNTR was tested extensively on data from different sequencing technologies, including Illumina and PacBio.

Like other STR genotyping tools, adVNTR works best when reads span the VNTR. However, even with this limitation, there are (a) close to 100,000 VNTRs in the genic regions of human genome that can be spanned by Illumina reads; (b) indel detection is possible even when RU counting is not, for long VNTRs; (c) lower bounds on RU counts can separate some pathogenic cases from normal cases particularly when the normal VNTR length is shorter than the read length, while the pathogenic case is much longer (e.g. *CSTB*). Finally, dropping costs for long read sequencing (esp. PacBio, and Nanopore) will allow us to span and genotype over 158, 000 genic VNTRs.

The choice between short and long read technologies offers some trade-offs. Specifically, long reads allow for the targeted genotyping of a larger set of VNTRs (559,804), and are becoming increasingly cost-effective. However, the large numbers of indels in these technologies reduce the accuracy somewhat, and they are best used when there is a big difference between normal and pathogenic cases in terms of RU counts, or when the VNTRs are too long to be spanned by Illumina.

In contrast, short-read Illumina sequencing is increasingly used for Mendelian pipelines, and can be easily extended to include VNTR genotyping, with higher accuracy than PacBio. Also, the large number of VNTRs (458, 158) that can be spanned by Illumina reads makes it the technology of choice for association testing and population based studies.

In this research, we also provided initial results on genotyping frameshift errors in coding VNTRs, focusing on the easier case when all RUs have the same length. Future work will focus on extending the target VNTRs for RU counting and frameshift detection for VNTRs that are of medical interest, population genetics of VNTRs, and algorithmic strategies for speeding up VNTR discovery and genotyping.

## 4 Method

A VNTR sequence can be represented as *SR*_1_*R*_2_ … *R_u_P*, where *S* and *P* are the unique flanking regions, and *R_i_*(1 ≤ *i* ≤ *u*) correspond to the tandem repeats. For each *i,j, R_i_* is similar in sequence to *R_j_*, and the number of occurrences, *u*, is denoted as the *RU count*. We do not impose a length restriction on *S* and *P*, but assume that they are long enough to be unique in the genome. For genotyping a VNTR in a donor genome, we focus primarily on estimating the diploid RU counts (*u*_1_, *u*_2_). However, many (~ 10^3^) VNTRs occur in coding regions, and mutations, particularly frameshift causing indels, are also relevant. Our method, adVNTR, models the problems of RU counting and mutation detection using HMMs trained for each target VNTR. adVNTR requires a one-time training of models for each combination of a VNTR and sequencing technology, although the user has the option to retrain models. Once models are trained, it has three stages for genotyping: (i) Read recruitment; (ii) RU count estimation; and, (iii) variant (indel) detection. We describe the training procedure and the three modules below.

### HMM Training

The goal of training is to estimate model parameters for each VNTR and each sequencing technology. Previous works have shown that an HMM with three groups of states could be used to find similarities between biological sequences (Eddy, 1996). In this model, a profile-HMMs can model a groups of sequences. Then, a new sequence can be aligned to a profile HMM to discover sequence family(Krogh et al., 1994). We use an HMM architecture with three parts, which have their own three groups of states (Fig. 5). The first part matches the 5’ (left) flanking region of the VNTR. The second part is an HMM which matches an arbitrary number of (approximately identical) repeating units. The last part matches the 3’ (right) flanking region (Fig. S1). The RU pattern is matched with a profile HMM (*RU HMM*), with states for matches, deletions, and insertions, and its model parameters are trained first. To train RU HMM for each VNTR, we collected RU sequences from the reference assembly(Lander et al., 2001) and performed a multiple sequence alignment(Eddy et al., 1995). Let *h*(*i,j*) denote the number of observed transitions from state *i* to state *j* in hidden path of each sequence in multiple alignment, and *h_i_*(*α*) denote the number of emissions of *α* in state *i*. We define permissible transition (arrows in Fig. 5) and match-state emission probabilities as follows:

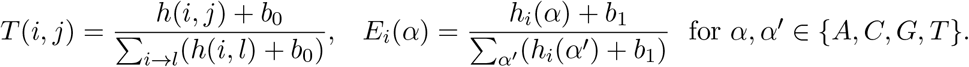

**Figure 5:**
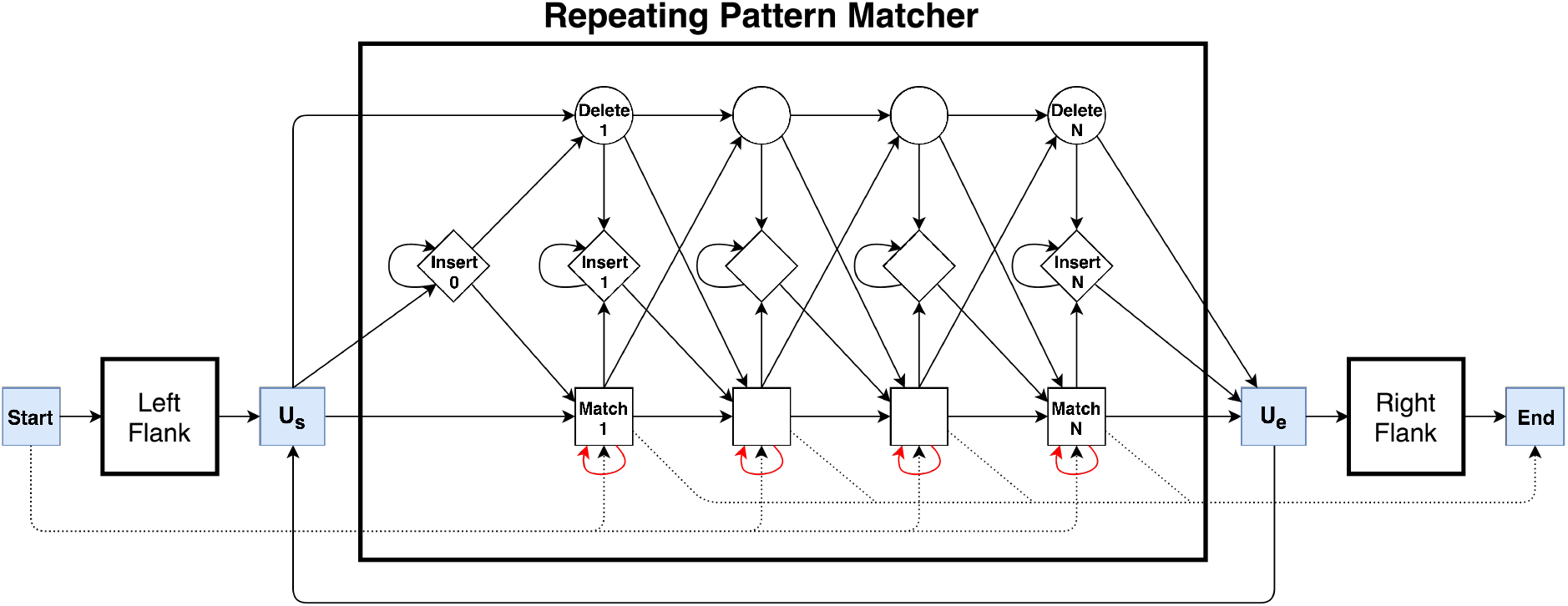
The VNTR HMM. The HMM is composed of 3 profile HMMs, one each for the left and right flanking unique regions, and one in the middle to match multiple and partial numbers of RUs. The special states *U_s_* (‘Unit-Start’), and *U_e_* (‘Unit-End’) are used for RU counting. Dotted lines refer to special transitions for partial reads that do not span the entire region.

Non-permissible transitions have probability 0, and *h_i_*(*α*) = 1/4 for insert state *i* and 0 for deletions. The pseudocounts *b*_0_ and *b*_1_ were estimated by initially setting them to the error rate of the sequencing technology, but they (along with other model parameters) were updated after aligning Illumina or PacBio reads to the model. The RU HMM architecture was augmented by adding (a) transitions from *U_e_* to *U_s_* to allow matching of variable number of RU; (b) adding the HMMs for the matching of any portions of left and right flanking sequences; and (c) by adding transitions to match reads that match either the left flanking or the right flanking region. In addition, reads anchored to one of the unique regions can jump past the other HMM using dotted arrows.

While error correction tools for PacBio have been developed, most do not work for repetitive regions,(Hackl et al., 2014; Salmela and Rivals, 2014; Au et al., 2012; Miclotte et al., 2016; Lee et al., 2014) and others assume a single haplotype for error correction(Salmela et al., 2016; Berlin et al., 2015). In contrast, the HMM allows us to model many of the common (homopolymer) errors directly. Insertion deletion errors are common in single molecule sequencing particularly in homopolymer runs of length ≥ 6, and occur mostly as insertions in the homopolymer run(Chaisson and Tesler, 2012). Consider a match state i with highest emission probability for nucleotide *α*. The transition probability *T*(*i,i*) from a match state *i* to itself was set based on the match probabilities of *α* in previous *k* = 6 states. The model parameters were further updated using genome sequencing data of NA12878 (Supplementary Material “Model Structure and Parameter Setting”).

### Read Recruitment

The first step in adVNTR is to *recruit* all reads that match a portion of the VNTR sequence. Alignment-based methods do not work well due to changes in RU counts (See Results), but the adVNTR HMM allows for variable RU count. To speed up recruitment, we used an Aho-Corasick keyword matching algorithm available as part of the Blast package(Altschul et al., 1990) to identify all reads that match a keyword from the VNTR patterns or the flanking regions. Note that the dictionary construction is a one-time process, and all reads must be scanned once for filtering. The keyword size and number of keywords were empirically chosen for each VNTR. Filtered reads were aligned to the HMM using the Viterbi algorithm. Only reads with matching probability higher than a specified threshold were retained. To compute the selection threshold for each VNTR, we aligned non-target genomic sequences that passed the keyword matching step to the HMM to form an empirical false distribution. Subsequently, we aligned VNTR encoding sequences to the HMM to form the score distribution of true reads. Then, we used a Naïve Bayes classifier to select a threshold.

### Estimating VNTR RU Counts

All reads covering an RU element are aligned, or ‘matched’ to the HMM using the Viterbi algorithm to create, in effect, a new multiple alignment. Recalling the Viterbi algorithm, let *V_k,j_* denote the highest (log) probability of emitting the first *k* letters of the sequence *s*_1_, *s*_2_,… *s_n_* and ending in state *j* of an HMM. Let, Prev_*k,j*_ denote the state *j*′ immediately prior to *j* in this optimum parse. Then,

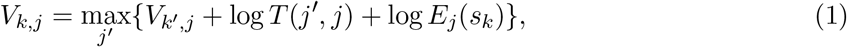

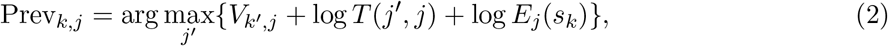

where, *k*′ = *k* − 1 for match or insert states; *k*′ = *k* otherwise.

For each read, the Viterbi algorithm allows for the enumeration of the maximum likelihood (ML) path by going backwards from Prev(End, *n*). Ignoring all but the *U_s_* and *U_e_* states in the Viterbi path, we get a pattern of the form 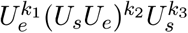 with *k*_1_, *k*_3_ ∈ {0,1}, and *k*_2_ ≥ 0. We estimate the RU count of the read as *k*_1_ + *k*_2_ + *k*_3_, and mark it as a lower bound if *k*_1_ + *k*_3_ > 0 (see Fig. 6 for an example).

**Figure 6:**
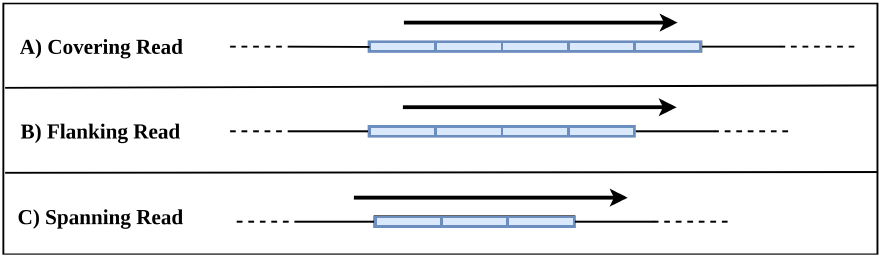
Estimates of RU counts using recruited reads. (A) (*k*_1_, *k*_2_, *k*_3_) = (1, 3,1); RU count ≥ 5. (B) (*k*_1_, *k*_2_, *k*_3_) = (0, 3,1); RU count ≥ 4 (C) (*k*_1_, *k*_2_, *k*_3_) = (0, 3, 0); RU count = 3.

One of the main reasons for erroneous RU counts is stutter during PCR amplification. The PCR amplification process is similar to replication errors that result on genetic RU count variation during cell-division, except that there are multiple rounds of amplification. In each PCR round, the number of copies might change by 1 with some probability. Once a single event has occurred and an erroneous template is generated, the event of having another change is likely to be independent of the previous event(Gymrek, 2016). To model errors in read counts, we define parameter *r_ϵ_* s.t. 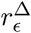 is the probability of RU counting error by ±Δ in the estimation of the true count. Thus the probability of getting the correct count is 1 − *r*, where

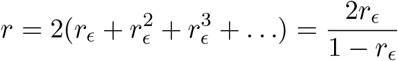

The analysis of reads at a VNTR gives us a multi-set of RU counts (or lower bounds) *c*_1_, *c*_2_,…,*c_n_*. We assume that the donor genome is diploid but do not require any phasing information in the computation of the multi-set. Additionally, we allow the possibility that all reads are sampled from one haplotype with the RU count of the missing haplotype being *X*. We define *C* = {*c*_1_, *c*_2_,…, *c_n_*} ∪ {*X*} and use *C* to get a list of possible genotypes (*c_i_,c_j_*) with *c_i_* ≤ *c_j_*. Then, the conditional likelihood of a read with RU count *c* is given by:

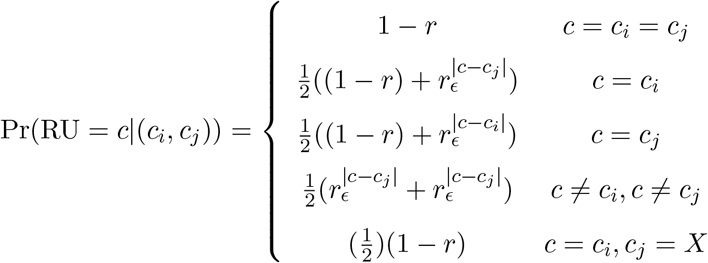

Similarly, the likelihood of a read with a lower bound *c* on the RU count is given by:

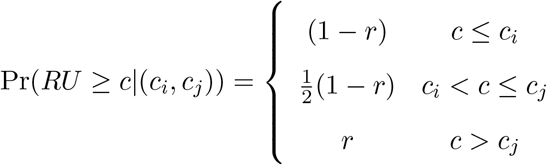

The likelihood of the data *C* is given by ∏_*c_k_*∈*C*_ Pr(*c_k_*|(*c_i_, c_j_*)). The posterior genotype probabilities can be computed using Bayes’ theorem:

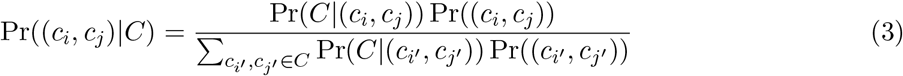

We generally set equal priors. However, in the event that we only see reads with a single count *c*’, we choose 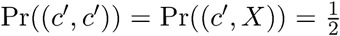. The probability of “missing haplotype” event is modeled as a Bernoulli process since in genome sequencing, sampling from either chromosome is done at random and so, the probability of not observing a halplotype in each read (failure) is 1/2. If we see multiple counts, we set Pr((*c*’,*X*)) = 0 for all *c*’ ∈ *C*, and give equal priors to all other genotypes.

### VNTR Mutation Detection

It is not difficult to see that alignment based methods do not work well in VNTRs. Changes in RU counts make it difficult to align reads even for mappers that allow split-reads, as the gaps in different reads can be placed in different locations. A similar problem appears with small indels, as there are multiple ways to align reads with an indel in a Repeat Unit. The adVNTR HMM aligns all repeat units to the same HMM, and this has the effect of aligning all mutations/indels in the same column. Consider the case where reads contain a total of *υ* nucleotides matching a VNTR RU of length *ℓ*, and RU count *u*. Moreover at a specific position covered by *d* Repeats, suppose we observe *ι* indel transitions.

For a true indel mutation, we expect 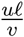 fraction of transitions at a location to be an indel, giving a likelihood of the observed data as Binom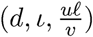. Alternatively, for a homopolymer run of *i* > 0 nucleotides, let *ε_i_* denote the per-nucleotide indel error rate. We modeled *ε*_1_ empirically in non-VNTR, non-polymorphic regions and confirmed prior results that *ε_i_* increases with increasing i(Margulies et al., 2005). Thus, the likelihood of seeing *ι* indel transitions due to sequencing error in a homopolymer run of length *i* is Binom(*d, ι,ε_i_*). We scored an indel in the VNTR using the log-likelihood ratio

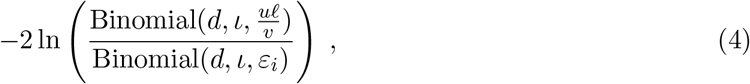

which follows a *χ*^2^ distribution. We select the indel if the nominal *p*-value is lower than 0.01.

Command line usage of adVNTR for RU count genotyping and frameshift identification is available in Supplementary Material “Running adVNTR”

## Acknowledgements

The analyses presented in this paper are based on the use of study data downloaded from the dbGaP web site, under phs001095.v1.p1, phs001096.v1.p1 and phs001097.v1.p1.

## Supplementary Material

### A. Model Structure and Parameter Setting

Each VNTR is represented by three Hidden Markov Models. A detailed sketch of the Repeat Match HMM is shown in Fig. 5. Here, we show the structure of two other parts in Fig. S1. We repeated the blue silent states (*Start, U_S_, U_e_*, and *End*) to show how these three models are connected.

To set the transition and emission probabilities of repeat matcher, we used the parameter obtained by pair HMM of repeating units in reference genome. We set pseudocounts equal to error rate of sequencing technology in all three HMMs to allow for mutations and sequencing errors. After the initialization of each model, we updated them using sequencing data of NA12878 (Table S2). To update each model, we ran read recruitment on sequencing data of NA12878 and extracted repeating units as described in Methods. Then, we aligned the repeating units to the HMM, and used the new aligned reads to update HMM parameters. We measure fitness of model by the sum of log-likelihood of the recruited reads, as follows:

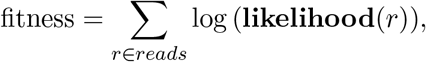

where likelihood of read *r* is defined as the probability of most likely path in the HMM to emit *r*. We continued to iterate the model alignment, and parameter update steps until convergence of fitness values.

As described in Methods, we compute the likelihood using the Viterbi algorithm. Let *V_k,j_* denote the highest (log) probability of emitting the first *k* letters of the sequence *s*_1_, *s*_2_, … *s_n_* and ending in state *j* of an HMM. Let, Prev_*k,j*_ denote the state *j*’ immediately prior to *j* in this optimum parse. Then,

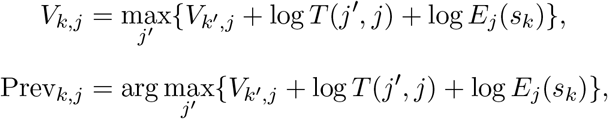

where, *k*’ = *k* − 1 for match or insert states; *k*’ = *k* otherwise. Then, for a read sequence *r* with length *n, max_j_V_n,j_* over all states *j* in the HMM determines the maximum likelihood.

**Figure S1:**
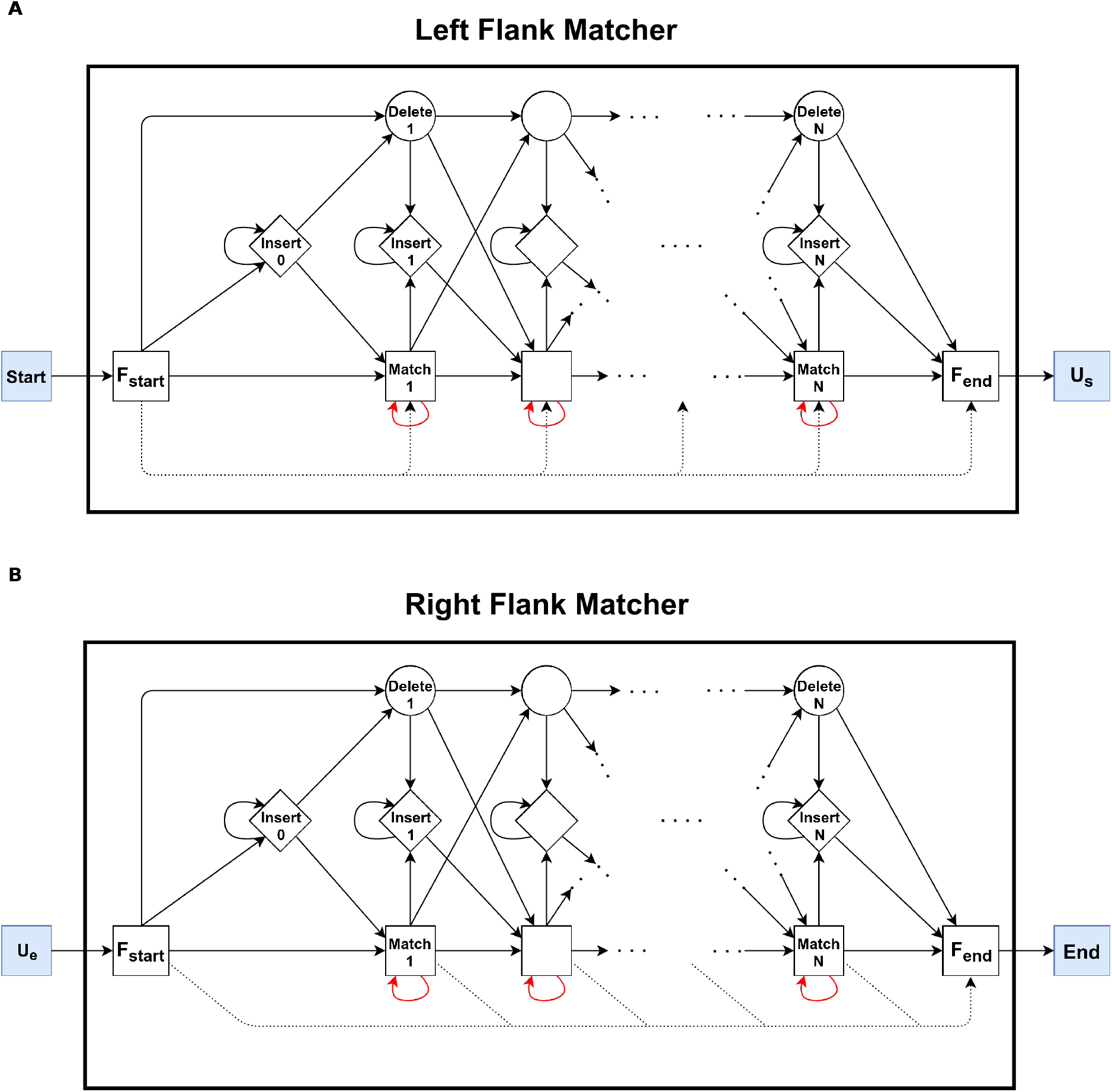
Flanking region matchers. (A) Shows the structure of Left Flank Matcher, which matches a suffix of left flanking region of the VNTR. In this part, the dotted edges allows skipping of adjacent base pairs at the beginning of the flanking region, and the rest of region (base pairs on the right) should be matched to the states and this is how matching of a suffix is insured. (B) Shows the structure of Right Flank Matcher, the model that matches a prefix of right flanking region of the VNTR. Here, dotted edges ensure the matching of a prefix of the flanking region sequence.

### B. Selecting Target VNTRs

We selected sets of target models that could be analyzed based on their characteristics and the sequencing technologies as follows: We started with the human VNTR list created by Tandem Repeat Finder. To select the most important loci, we considered VNTRs that had an intersection with coding regions of human genome. Next, we excluded cases where the flanking regions of VNTR were not known (e.g. VNTR is close to telomere; the flanking region doesn’t exist in reference genome; and there is a sequence of ‘N’ adjacent to the VNTR.). Finally, we added 17 VNTRs that are in promoter or intron of the genes but are known to be linked to a disease (Table 1). We removed VNTRs that appear multiple times in different loci of the genome with identical patterns and flanking regions, but with different number of copies. To find such similar VNTRs, we compared each pair of VNTRs by comparing the flanking regions and repeating unit with BLAT (Kent, 2002) and eliminating the VNTRs if their similarity was higher than 75%.

This procedure resulted in 2944 ‘coding’ VNTRs out of 3147 VNTRs that intersected with coding regions of human genome. The 2944 VNTRs were used for PacBio analysis. For Illumia analysis, we used a subset of 1775 VNTRs of the 2944, whose length was shorter than 140bp. Finally to create a difficult test case for testing frame-shifts, we selected 115 of 2944 VNTRs for which the total length was ≥ 250bp, and all Repeat Units had the same length, and used those to simulate indel (frameshift) data-sets.

### C. Test Datasets

Multiple test cases were generated using the three lists containing 2944, 1775, and 115 VNTRs, respectively as described in the previous section. We started by generating a distinct human genomic sequence `VNTR_I_X_reference.fa` for each *I* ∈ [1, 2944] and each value *X* ∈ [−3, 3] (20,608 total sequences). Each sequence `VNTR_I_X_`reference.fa was identical to the human reference except that it had X’ copies for **I**-th `VNTR`, where X’ takes the RU count in reference genome ±*X*. To increase the `RU` count of a `VNTR`, we added the repeating units from the first repeat to the last unit, one at a time. We additionally generated ~ 4920 reference sequences `VNTR_I_Deletion_P.fa` and `VNTR_I_Insertion_P.fa` for all *I* ∈ [1,115] `VNTRs` indexing the third list, and a single insertion or deletion at the *P*th base pair of the *I*th `VNTR`. We set *P* to every position in the VNTR that was a multiple of 10 and was at least 140bp apart from each side of the VNTR. These reference templates were used for generating simulated datasets as follows:

#### IlluminaSim Dataset

We used the following command to simulate the reads from haplotypes using ART:

~~~
art_illumina -ss HSXt -sam -i VNTR_I_X_reference.fa -l 150 -f 15 -o VNTR_I_X_set
~~~

Then, we merged every pair of haploid datasets with RU counts X and Y to get diploid sequencing data with genotype (X,Y) for `VNTR` *I* by appending `VNTR_I_X_`set.fq to the end of `VNTR_I_Y_`set.fq to get `VNTR_I_XY_`set.fq. Then, we aligned these diploid reads to the reference genome using Bowtie2 as follows:

~~~
bowtie2 -x hg19_bowtie2_index -U VNTR_I_XY_set.fq -S VNTR_I_XY_aln.sam
~~~

#### PacBioSim Dataset

We used the following command to simulated the reads for *I*th VNTR using SimLoRD:

~~~
simlord -rr VNTR_I_X_reference.fa -pi 0.12 -pd 0.02 -ps 0.02 -c 15 VNTR_I_X_pb_set.sam
~~~

Next, we merged each pair of reads (fastq files) to get the diploid set of reads at 30X coverage.

#### PacBioLong Dataset

The dataset is similar to PacBioSim but with higher RU counts for 3 VNTRs 120, 40, and 25 for VNTRs in *INS, CSTB*, and *HIC1* genes, which represent the largest expansion known for these VNTRs. Again, we used SimLord to generate reads.

~~~
simlord -rr VNTR_I_X_reference.fa -pi 0.12 -pd 0.02 -ps 0.02 -c 30 VNTR_I_X_pb_set.sam
~~~

#### PacBio Coverage Dataset

We simulated different levels of coverage for the three VNTRs using:

~~~
simlord -rr VNTR_I_X_reference.fa -pi 0.12 -pd 0.02 -ps 0.02 -c C VNTR_I_X_C_set
~~~

Here, 1 ≤ *C* ≤ 40 ×.

#### IlluminaFrameshift Dataset

We simulated these datasets using following commands:

~~~
art_illumina -ss HSXt -sam -i VNTR_I_Deletion_P.fa -l 150 -f 15 -o VNTR_I_Deletion_p
art_illumina -ss HSXt -sam -i VNTR_I_Insertion_P.fa -l 150 -f 15 -o VNTR_I_Insertion_p
~~~

We also simulated reads from reference genome without the frameshift:

~~~
art_illumina -ss HSXt -sam -i hg19.fa -l 150 -f 15 -o normal_haplotype
~~~

Finally, we merged fastq read files of a haplotype with frameshift with that of normal haplotype to get the diploid sample at 30X coverage and aligned the reads with Bowtie2 similar to “IlluminaSim Dataset”.

**Table S1:** Simulated dataset summary.

| Dataset Name | Profile | Depth | # of VNTRs |
| --- | --- | --- | --- |
| Illumina Genotyping Dataset | HiSeqX TruSeq | 30X | 1775 |
| PacBio Genotyping Dataset | PacBio | 30X | 2944 |
| PacBio Long Expansion Dataset | PacBio | 30X | 3 |
| PacBio Coverage Dataset | PacBio | $1 \leq C \leq 40$ | 3 |
| Frameshift Dataset | HiSeqX TruSeq | $C \in \{10, 20, 30, 40\}$ | 123 |

WGS data used for testing was taken from Genome in a Bottle, NCBI short read archive, Polaris, while exome data was obtained from GoT2D. See Table S2

**Table S2:** Real sequencing data used in tests.

| Samples | Study | Profile | PCR free | Depth | Access |
| --- | --- | --- | --- | --- | --- |
| AJ Child (NA24385) | GIAB | PacBio | - | 70X | http://jimb.stanford.edu/giab-resources |
| AJ Father (NA24149) | GIAB | PacBio | - | 30X | http://jimb.stanford.edu/giab-resources |
| AJ Mother (NA24143) | GIAB | PacBio | - | 30X | http://jimb.stanford.edu/giab-resources |
| Chinese Child (HG00514) | PRJEB12236 | PacBio | - | 70X | ncbi.nlm.nih.gov/sra/ERX1322863 |
| Chinese Father (HG00512) | PRJEB12236 | PacBio | - | 35X | ncbi.nlm.nih.gov/sra/ERX1322861 |
| Chinese Mother (HG00513) | PRJEB12236 | PacBio | - | 35X | ncbi.nlm.nih.gov/sra/ERX1322862 |
| AJ Child (NA24385) | GIAB | HiSeq 2500 | Y | 40X | http://jimb.stanford.edu/giab-resources |
| AJ Father (NA24149) | GIAB | HiSeq 2500 | Y | 40X | http://jimb.stanford.edu/giab-resources |
| AJ Mother (NA24143) | GIAB | HiSeq 2500 | Y | 40X | http://jimb.stanford.edu/giab-resources |
| NA12878 | GIAB | PacBio | - | 70X | http://jimb.stanford.edu/giab-resources |
| NA12878 | GIAB | HiSeq 2500 | Y | 30X | http://jimb.stanford.edu/giab-resources |
| 150 Individuals of 1KGP | Polaris | HiSeq X | Y | 30-40X | ebi.ac.uk/ena/data/view/PRJEB20654 |
| Diabetes WES data | GoT2D | HiSeq 2000 | N | 82X | dbGaP: phs001095, phs001096, and phs001097 |

### D. Running adVNTR

adVNTR is available at https://github.com/mehrdadbakhtiari/adVNTR. As stated in the repository, the best way to install it is to use conda package manager and running `conda install advntr`. After installation, `advntr` command invokes the program with four possible commands `genotype, addmodel, viewmodel`, and `delmodel`. Detail of each command as well as complete tutorial on installation and usage are available at http://advntr.readthedocs.io/. Also, passing `-h` argument to each command will show the correct command line usage of the command.

### E. VNTRseek

In order to make a call on a VNTR, VNTRseek requires both ends to be anchored with a minimum of 20bp on each side of VNTR. This limits the length of VNTRs that can be identified using VNTRseek is limited to 110bp using Illumina sequencing technology. Also, it compares each VNTR in the sequencing reads to every VNTR in reference genome which makes the process computationally demanding, and inaccessible for large data-sets. For these reasons, extensive VNTRseek comparisons were not conducted.

### F. Supplementary Figures and Tables

**Figure S2:**
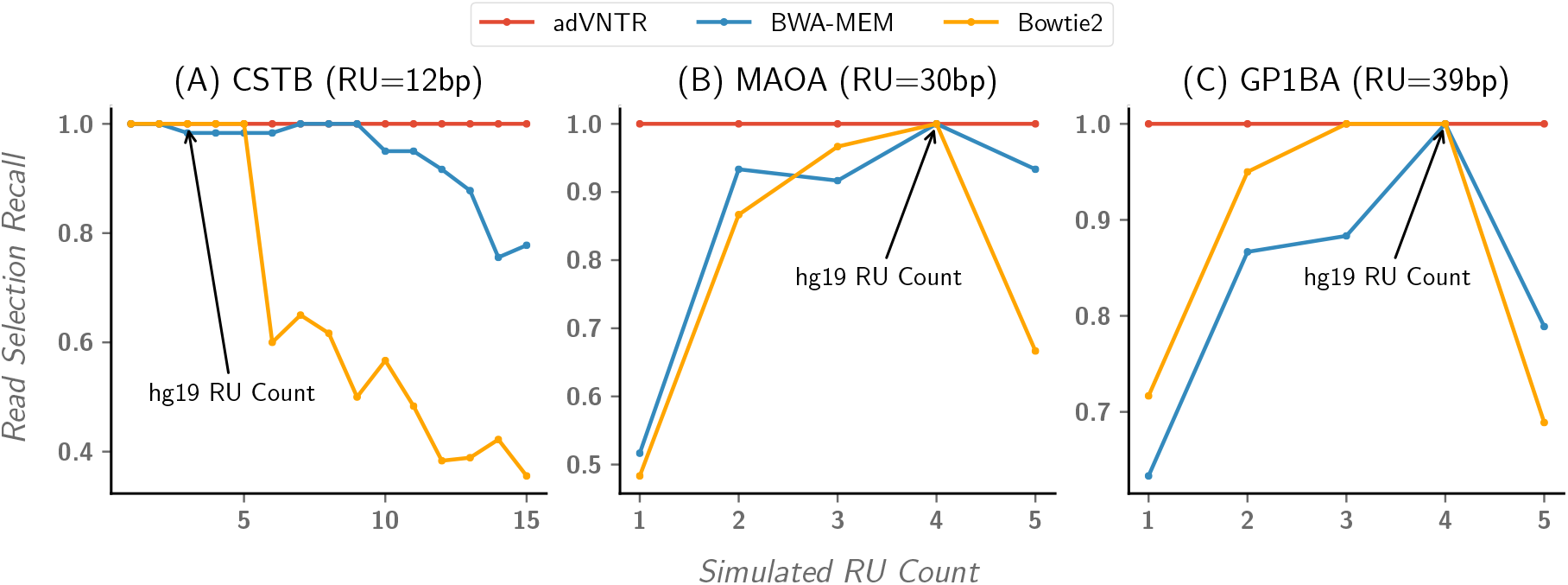
Sensitivity of Illumina read recruitment at specific VNTR loci. Comparison of adVNTR read selection with BWA-MEM and Bowtie2 mapping for Illumina reads (short VNTRs). Each plot shows the sensitivity of mapped/selected reads as a function of the number of repeats for different VNTRs. These plots show examples of alignment tools’ behavior when RU count of VNTR deviates from the RU count in the reference genome. (A) Shows the comparison for the VNTR in *CSTB* gene, in which the pathogenic cases have more then 12 repeats and as it is shown alignment tools perform poorly in those cases. (B) Shows the comparison for the VNTR in MAOA gene, where the 4 repeats corresponds to both pathogenic case and number of repeats in reference genome. However, other tools perform poorly in normal cases. (C) Shows the comparison for the VNTR in *GP1BA* gene, and again, alignment tools only perform well when RU count is same as RU count in reference genome.

**Figure S3:**
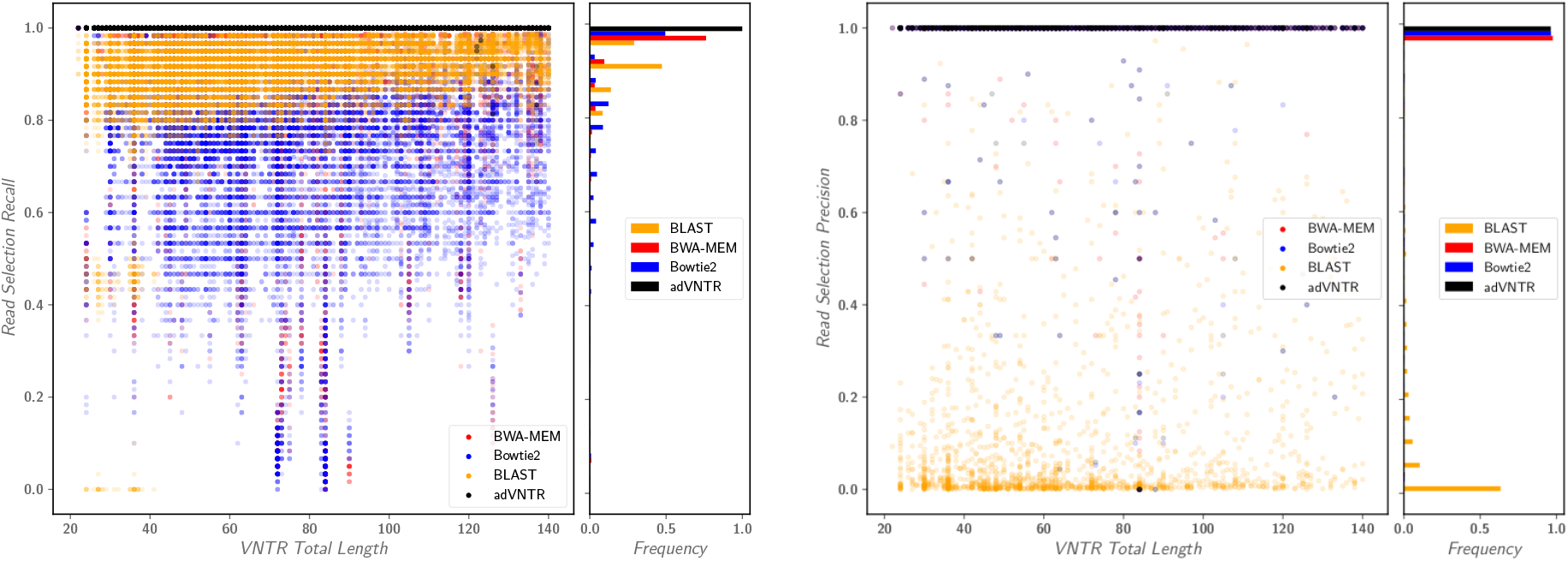
Read recruitment quality on Illumina reads. (A) Shows the comparison of the recall of adVNTR read recruitment with BWA-MEM, Bowtie2, and BLAST. (B) Shows the precision for read recruitment. These figures show that adVNTR has much higher recall compare to standard alignment tools without losing precision.

**Figure S4:**
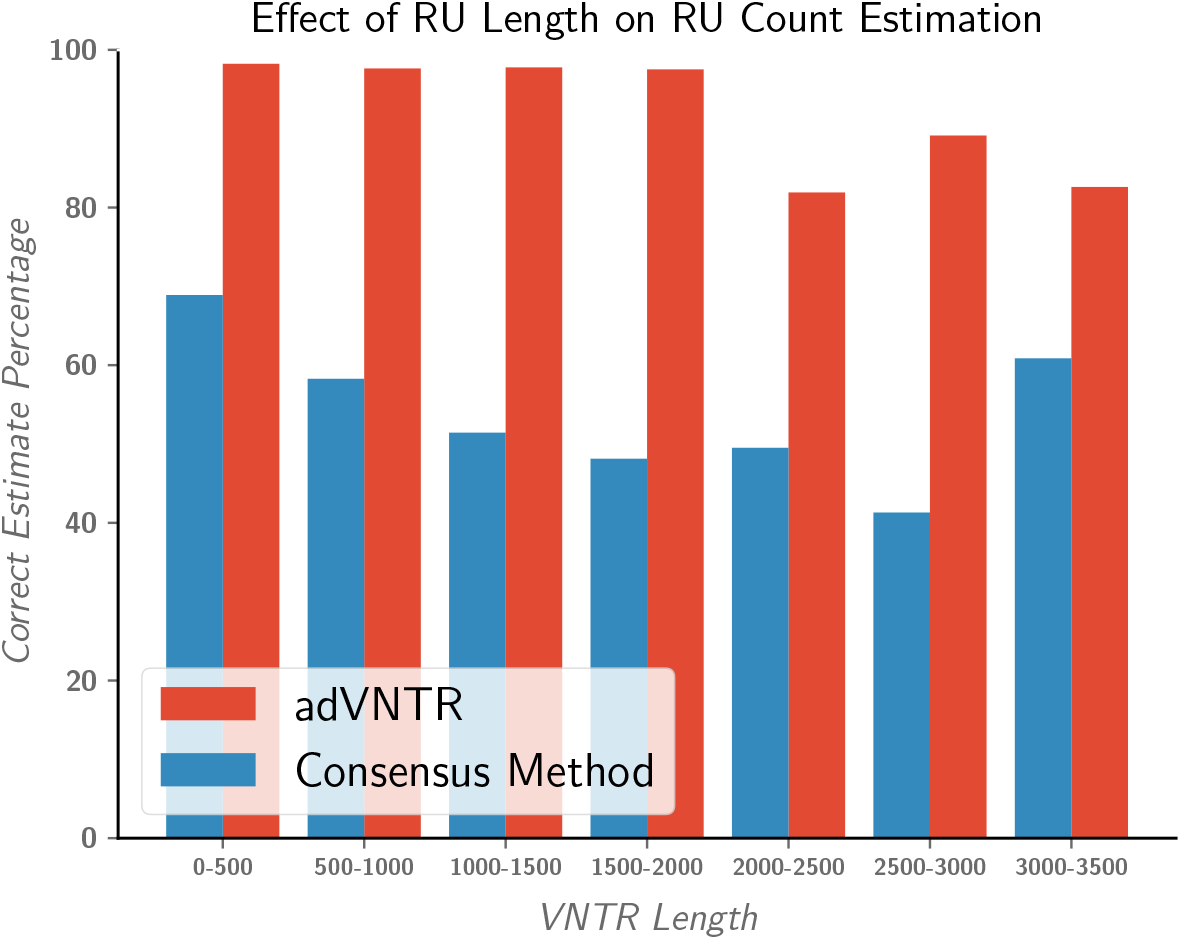
Comparison of adVNTR genotyping with consensus method on homozygous simulated data. adVNTR and consensus method comparison on homozygous testcases in *PacBioSim*.

**Figure S5:**
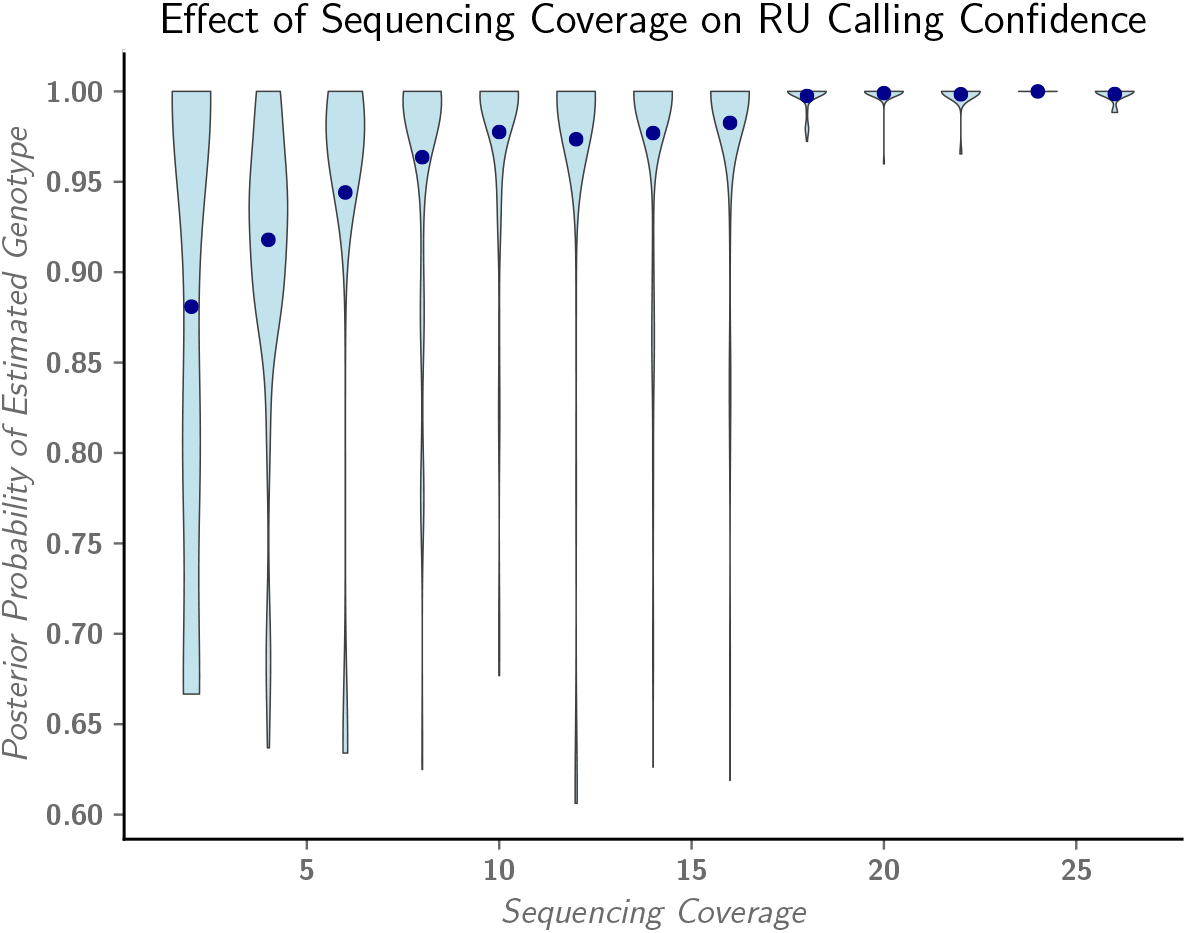
Association of PacBio sequencing coverage in VNTR region and posterior probability of RU count calling. The figure shows posterior probability of RU count estimation in AJ trio sequencing data form GIAB. Most of calls with low posterior probability (low confidence calls) result from low coverage in VNTR region. With at least 10 reads that span the VNTR, we will get 0.98 posterior probability for estimated genotype.

**Figure S6:**
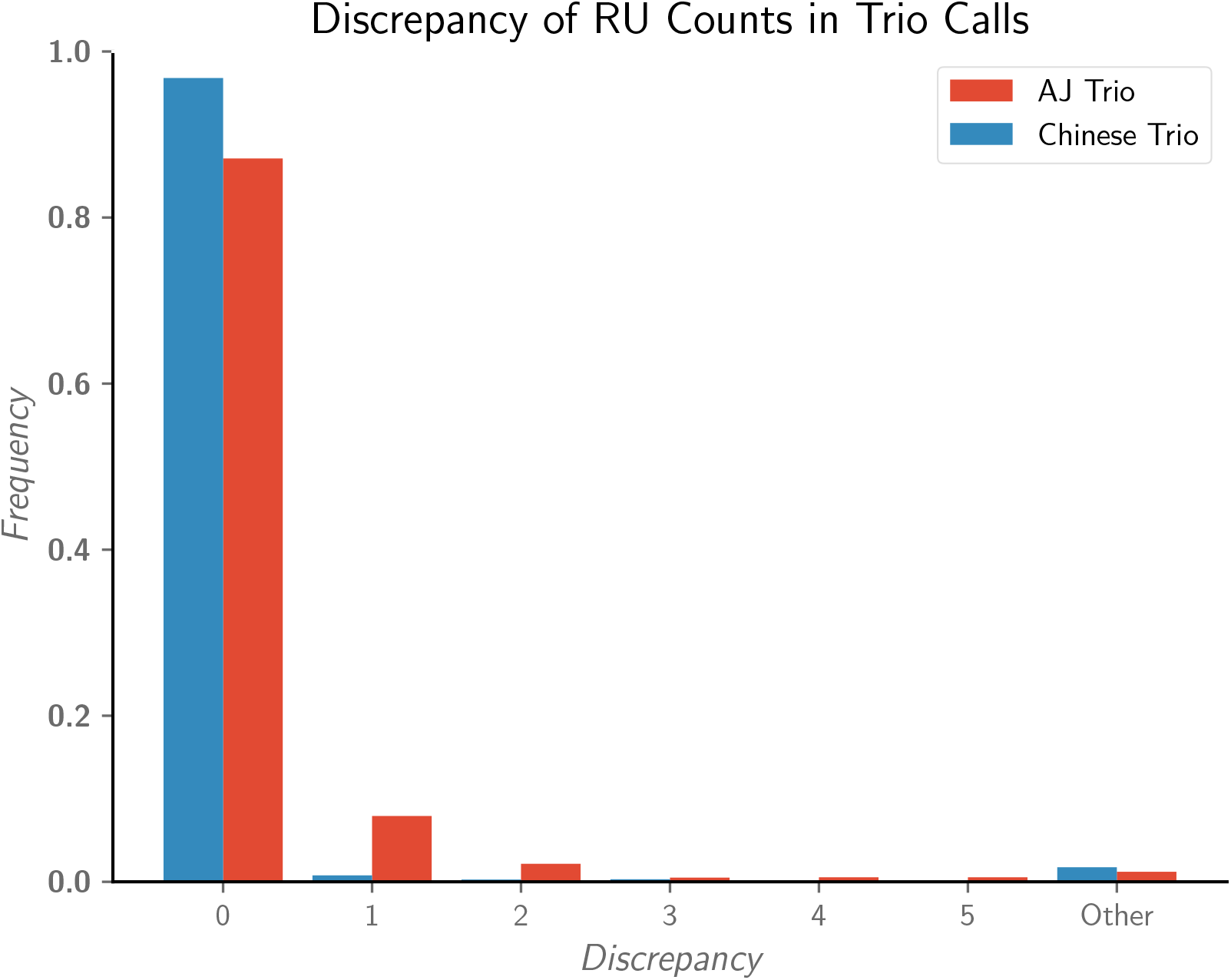
Distribution of discrepancies on trio calls using PacBio reads. This figure shows the distribution of discrepancies in adVNTR estimates on AJ and Chinese trios. As shown in the figure, most of non consistent calls in AJ trio have one discrepancy in estimated RU counts.

**Table S3:**
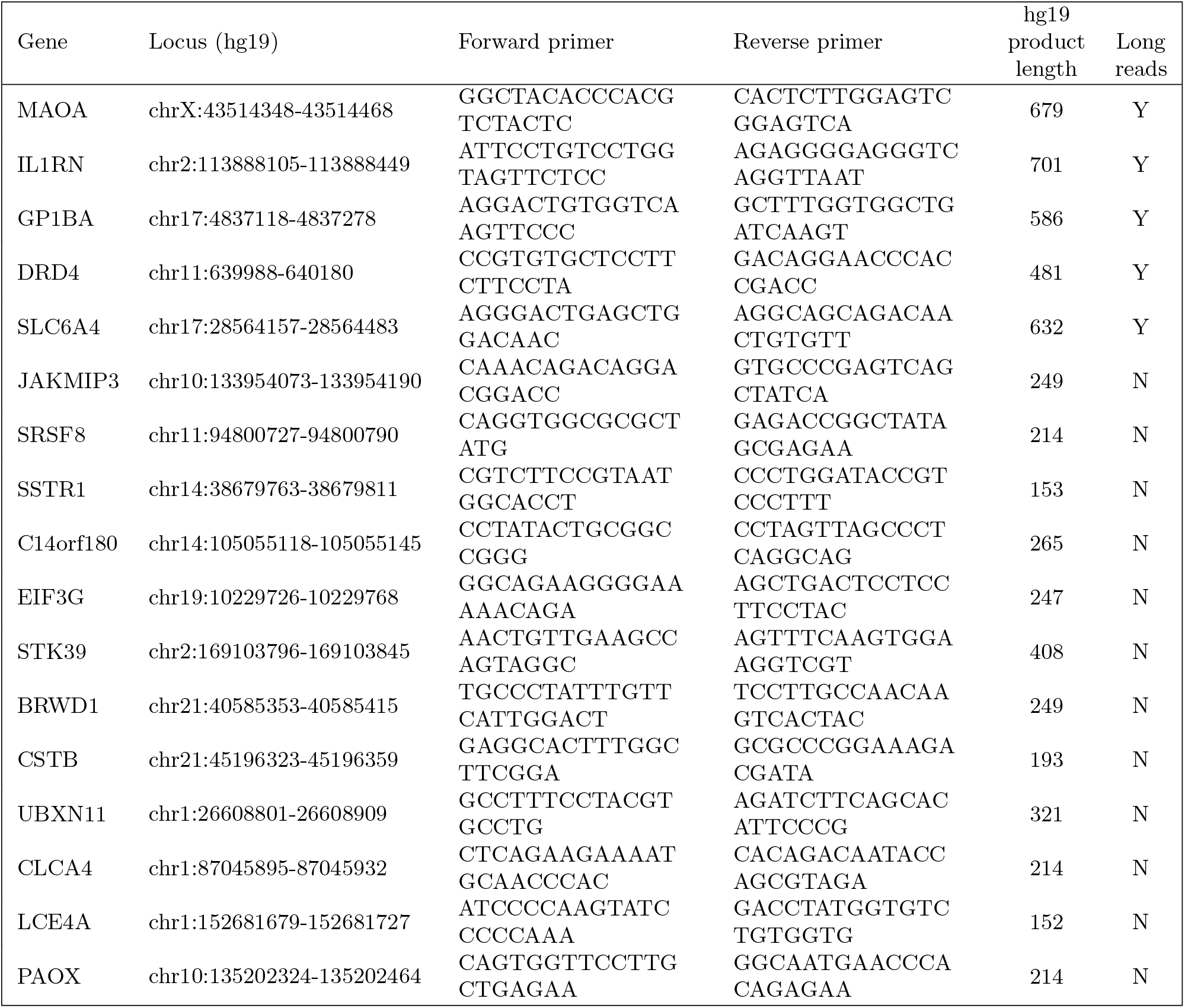
Primer for gel electrophoresis. Last column shows whether we used the primers were used for a long range PCR. We used long range PCR to validate adVNTR calls on longer VNTRs (using PacBio reads).

**Figure S7:**
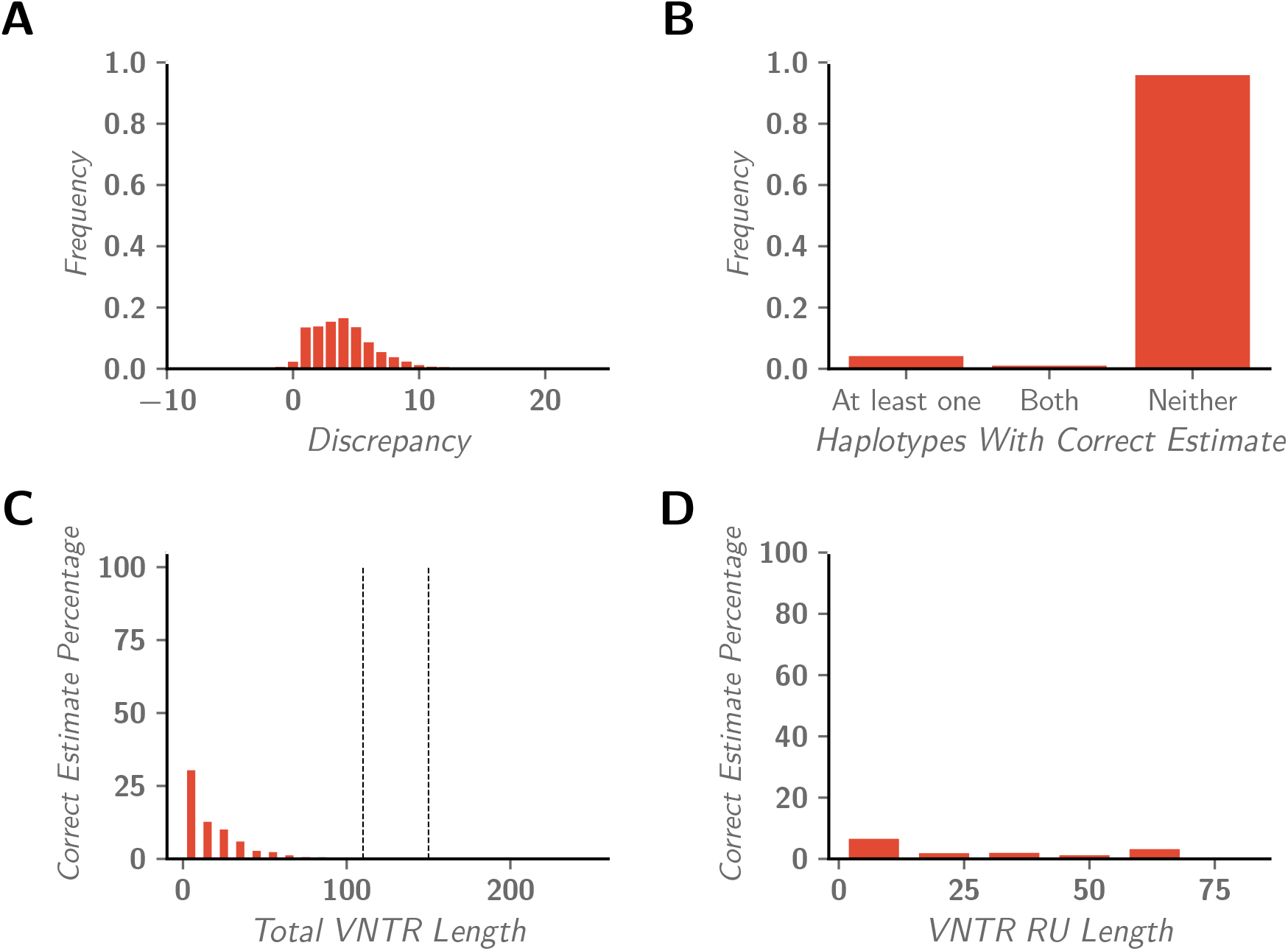
Expansion Hunter’s performance on VNTR genotyping using Illumina reads. Expansion Hunter’s performance on IlluminaSim dataset.

**Figure S8:**
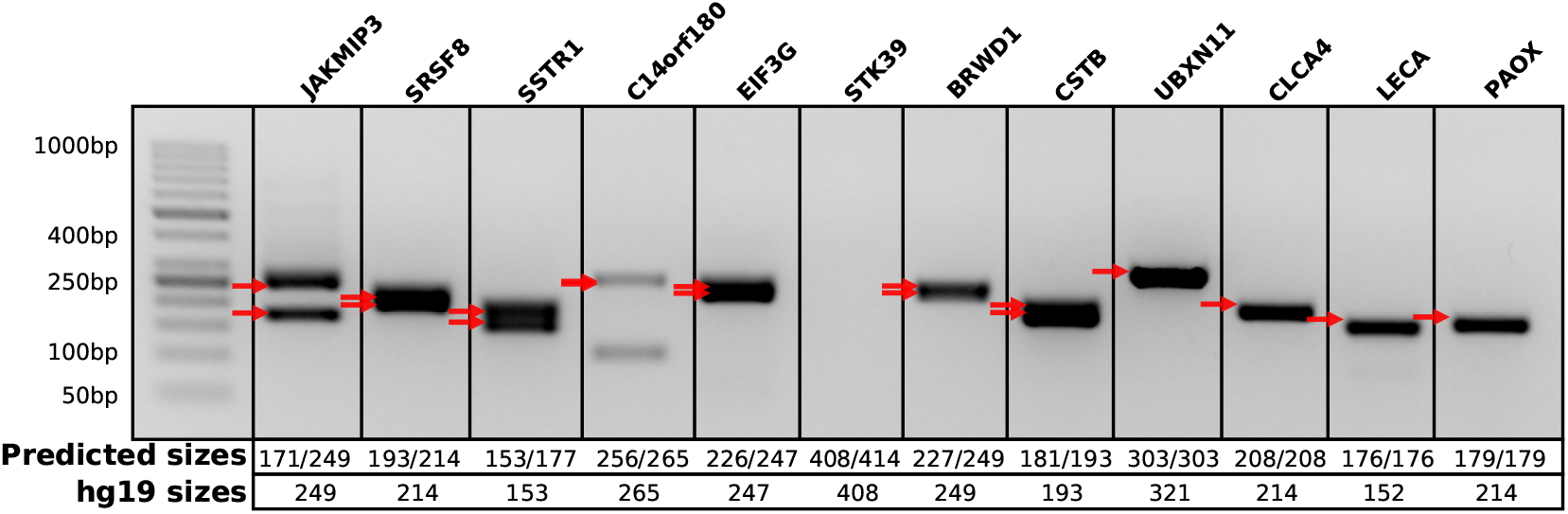
Validation of adVNTR genotyping on short VNTRs. In experiment for *C14orf180* the primers were repeated in another region of genome which resulted in having extra band. Even with zero copy of VNTR patterns, the distance of primers around VNTR is 238bp which means the extra band (~100bp) is resulted from another region of genome. Also, PCR amplification failed for *STK39* and no band is visible. Results of all other 10 experiments are consistent with adVNTR’s estimates.

**Table S4:** RU count genotyping results on simulated data. For two cases, (MAOA 1/1 and CSTB 1/1) Expansion Hunter doesn’t find any RU count.

| VNTR | Simulated Genotype | RU Count Discrepancy |  |  |
| --- | --- | --- | --- | --- |
|  |  | PacBio Dataset | Illumina Dataset |  |
|  |  | adVNTR | Expansion Hunter | adVNTR |
| MAOA | 1/1 | 0/0 | -/- | 0/0 |
| MAOA | 1/2 | 0/0 | 0/-1 | 0/0 |
| MAOA | 1/3 | 0/0 | 0/-2 | 0/0 |
| MAOA | 1/4 | 0/0 | 0/-3 | 0/0 |
| MAOA | 1/5 | 0/0 | 0/-4 | 0/0 |
| MAOA | 2/2 | 0/0 | -1/-1 | 0/0 |
| MAOA | 2/3 | 0/0 | 0/-1 | 0/0 |
| MAOA | 2/4 | 0/0 | -1/-3 | 0/0 |
| MAOA | 2/5 | 0/0 | -1/-4 | 0/0 |
| MAOA | 3/3 | 0/0 | -2/-2 | 0/0 |
| MAOA | 3/4 | 0/0 | -2/-3 | 0/0 |
| MAOA | 3/5 | 0/0 | -2/-4 | 0/0 |
| MAOA | 4/4 | 0/0 | -3/-3 | 0/0 |
| MAOA | 4/5 | 0/0 | -3/-4 | 0/0 |
| MAOA | 5/5 | 0/0 | -4/-4 | -1/-1 |
| GP1BA | 1/1 | 0/0 | 0/0 | 0/0 |
| GP1BA | 1/2 | 0/0 | 0/0 | 0/0 |
| GP1BA | 1/3 | 0/0 | 0/-1 | 0/0 |
| GP1BA | 1/4 | 0/0 | 1/-2 | 0/-1 |
| GP1BA | 2/2 | 0/0 | 0/0 | 0/0 |
| GP1BA | 2/3 | 0/0 | 0/-1 | 0/0 |
| GP1BA | 2/4 | 0/0 | 0/-2 | 0/-1 |
| GP1BA | 3/3 | 0/0 | -1/-1 | 0/0 |
| GP1BA | 3/4 | 0/0 | -1/-2 | 0/0 |
| GP1BA | 4/4 | 0/0 | -2/-2 | -1/0 |
| CSTB | 1/1 | 0/0 | -/- | 0/0 |
| CSTB | 1/2 | 0/0 | 1/0 | 0/0 |
| CSTB | 1/3 | 0/0 | 2/0 | 0/0 |
| CSTB | 1/4 | 0/0 | 3/0 | 0/0 |
| CSTB | 1/5 | 0/0 | 4/0 | 0/0 |
| CSTB | 1/6 | 0/0 | 4/-1 | 0/0 |
| CSTB | 1/7 | 0/0 | 3/-3 | 0/0 |
| CSTB | 1/8 | 0/0 | 4/-3 | 0/0 |
| CSTB | 1/9 | 0/0 | 3/-5 | 0/0 |
| CSTB | 1/10 | 0/0 | 4/-5 | 0/0 |
| CSTB | 1/11 | 0/0 | 4/-6 | 0/0 |
| CSTB | 1/12 | 0/0 | 4/-7 | 0/0 |
| CSTB | 1/13 | 0/0 | 4/-8 | 0/0 |
| CSTB | 1/14 | 0/0 | 3/-10 | 0/-1 |
| CSTB | 2/2 | 0/0 | 0/0 | 0/0 |
| CSTB | 2/3 | 0/0 | 1/0 | 0/0 |
| CSTB | 2/4 | 0/0 | 1/-1 | 0/0 |
| CSTB | 2/6 | 0/0 | 3/-1 | 0/0 |
| CSTB | 2/8 | 0/0 | 3/-3 | 0/0 |
| CSTB | 2/10 | 0/0 | 3/-5 | 0/0 |
| CSTB | 2/12 | 0/0 | 3/-7 | 0/0 |
| CSTB | 2/14 | 0/0 | 3/-9 | 0/-1 |
| CSTB | 3/3 | 0/0 | -1/-1 | 0/0 |
| CSTB | 3/4 | 0/0 | 2/1 | 0/0 |
| CSTB | 3/6 | 0/0 | 2/-1 | 0/0 |
| CSTB | 3/8 | 0/0 | 2/-3 | 0/0 |
| CSTB | 3/10 | 0/0 | 2/-5 | 0/0 |

**Table S5:**
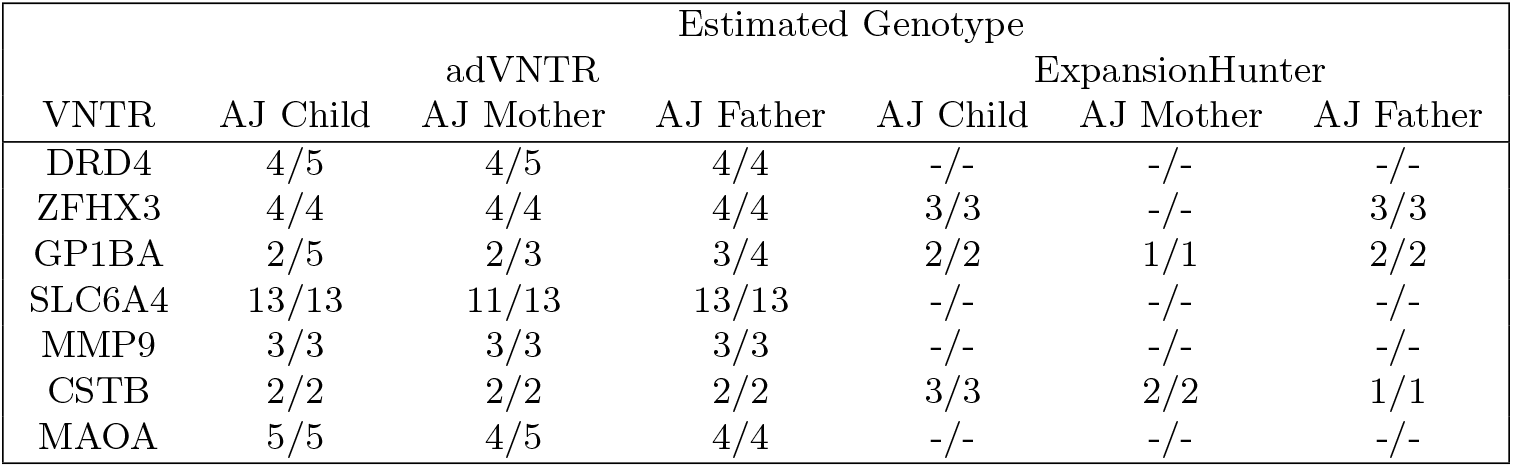
Genotyping comparison on AJ trio using Illumina reads from GIAB. Table shows the genotype found by adVNTR and ExpansionHunter in disease causing VNTRs that are shorter than Illumina reads. −/− denotes ExpansionHunter has not found any genotype for the VNTR. It worths mentioning the genotypes found by adVNTR for *MAOA* are not inconsistent as this VNTR is located on ChrX and the son has haploid RU counts inherited from mother.

**Figure S9:**
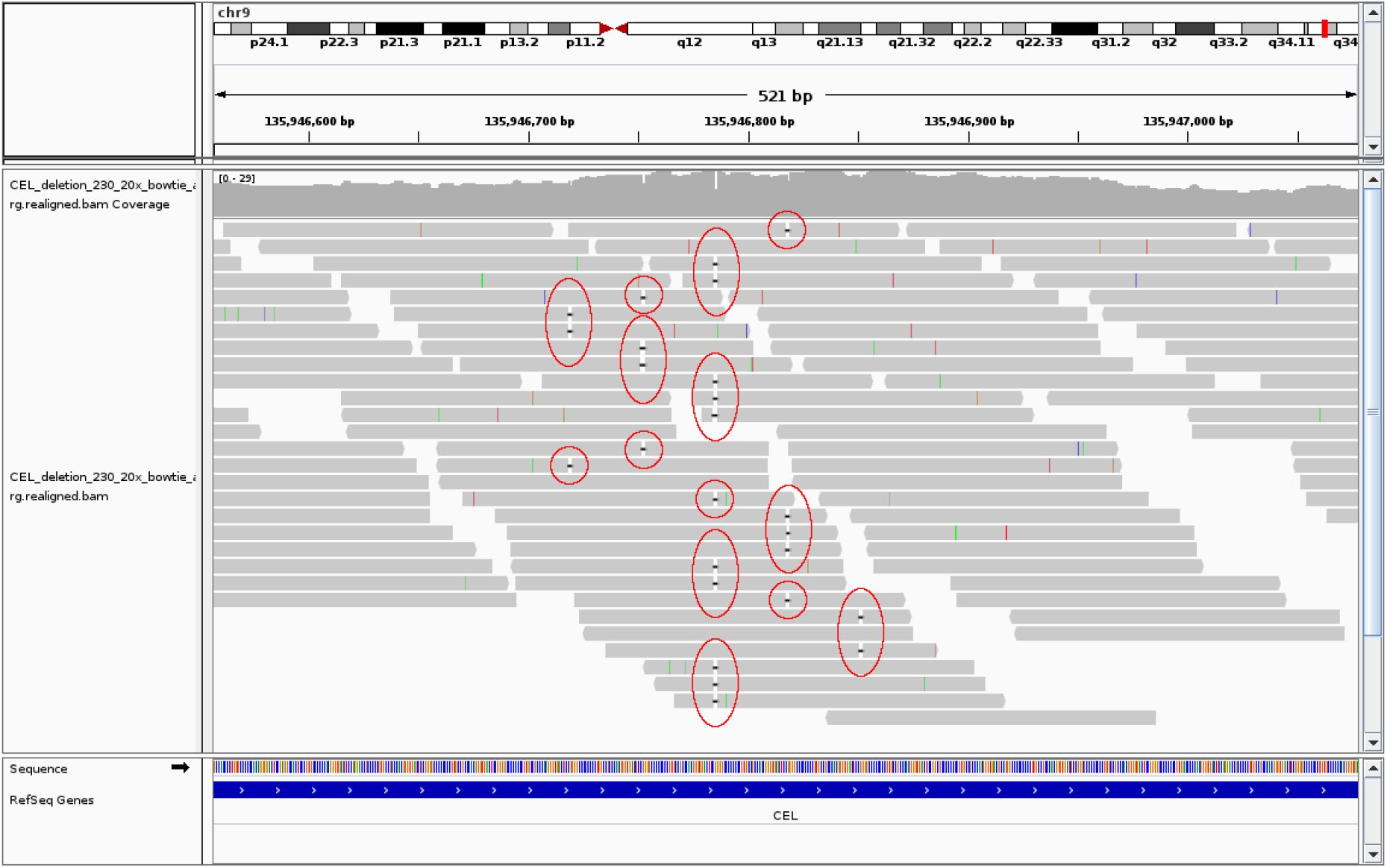
Alignment stats with frameshift. Alignment of a simulated data after running GATK IndelRealigner, when there is a deletion. With a sequencing mean of 30X, 25 reads contain the deletion but even after running realigner, deletions are mapped to five different repeating units.

**Figure S10:**
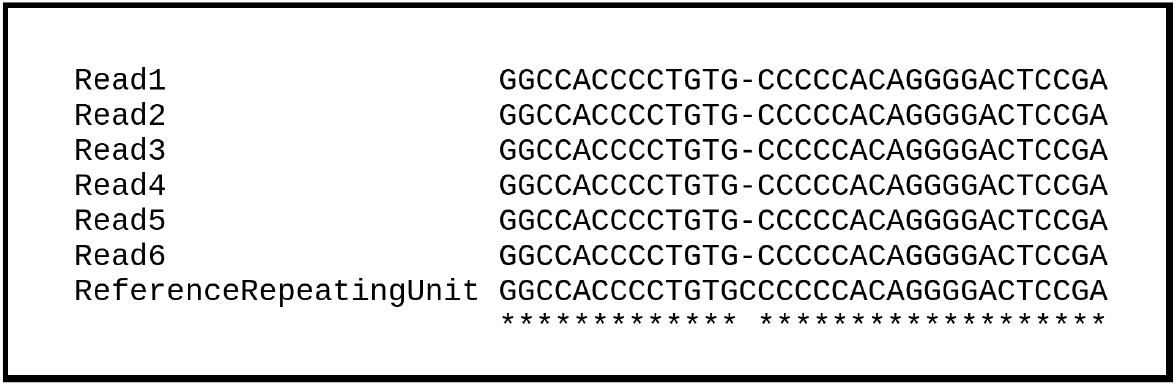
Frameshift in CEL gene. Multiple alignment of sequenced reads and reference repeating unit shows a deletion in diabetes patient genome. Due to low PCR amplification in GC rich VNTR region (84.8%), the coverage of VNTR region is 14X and 6 reads support the deletion.

**Table S6:** Comparison of indel finding with Samtools and GATK

|  |  | # of Samples | # of samples that frameshift has been identified |  |  |
| --- | --- | --- | --- | --- | --- |
|  |  |  | Samtools | Our Method | GATK |
| 10X | Insertions | 20 | 0 | 20 | 0 |
|  | Deletions | 20 | 0 | 20 | 0 |
| 20X | Insertions | 20 | 0 | 20 | 0 |
|  | Deletions | 20 | 0 | 20 | 0 |
| 30X | Insertions | 20 | 0 | 20 | 0 |
|  | Deletions | 20 | 0 | 20 | 0 |
| 40X | Insertions | 20 | 0 | 20 | 0 |
|  | Deletions | 20 | 0 | 20 | 0 |

